# Sex does not Influence Neuronal Autophagy throughout Aging in Mice

**DOI:** 10.1101/2025.01.16.633428

**Authors:** Mya N Rodriguez, Andrea KH Stavoe

## Abstract

Autophagy is critical for the homeostasis and function of neurons, as misregulation of autophagy has been implicated in age-related neurodegenerative diseases and neuron-specific knockdown of early autophagy genes results in early neurodegeneration in mice. We previously found that autophagosome formation decreases with age in murine neurons. Sex differences have been intensely studied in neurodegenerative diseases, but whether sex differences influence autophagy at the neuronal level have not been investigated. We compared protein expression of 22 autophagy components between neural tissues of female and male mice across development and aging. We found minimal sex-related differences in autophagy protein expression throughout the murine lifespan. Additionally, we assayed the recruitment of autophagy complexes and autophagosome biogenesis; we found no sex-dependent differences in multiple stages of autophagosome formation in neurons, independent of age. Our data suggest that biological sex does not influence autophagosome formation in neurons across development and aging.

## INTRODUCTION

Macroautophagy (hereafter autophagy) is a cellular homeostatic pathway that is especially critical to neuronal health and survival. Autophagy misregulation has been implicated in age-related neurodegenerative diseases, including Alzheimer, Parkinson, and Huntington Diseases (Menzies et al., 2017; Nixon, 2013; Yamamoto and Yue, 2014). Neuron-specific knock-out of autophagy components ATG5, ATG7, or EPG5 leads to early neurodegeneration in mice, highlighting the importance of autophagy in neurons (Hara et al., 2006; Komatsu et al., 2006; Zhao et al., 2013). Importantly, there are many instances of sex-related differences in diseases and disorders associated with autophagy misregulation, including neurodegenerative diseases (Shang et al., 2020). For example, females have higher risk of Alzheimer Disease, which is underpinned by sex differences in the regulation of autophagy (Congdon, 2018). Despite the clear importance of autophagy to neuronal function, autophagy has been primarily studied in yeast and mammalian immortalized cell lines, where the role of biological sex cannot be assessed. Furthermore, biological sex is often ignored in studies using primary mammalian neuronal cultures, leaving unknown whether biological sex influences neuronal autophagy at a cellular level.

Autophagosomes are formed by the concerted action of multiple protein complexes. The induction (or initiation) complex is composed of ULK1, ATG13, FIP200, and ATG101. ULK1 is a serine/threonine kinase that phosphorylates several other autophagy components. The nucleation complex generates PI3P and is composed of ATG14, Beclin1, PIK3C3/VPS34, and PIK3R4/VPS15. ATG14 and Beclin1 are members of the nucleation complex that are specific to autophagy; the other nucleation complex members also function in other cellular processes. The elongation complex consists of two ubiquitin-like conjugation complexes and has several protein constituents: ATG3, ATG4, ATG5, ATG7, ATG10, ATG12, ATG16L1. The elongation complex ultimately functions to conjugate ubiquitin-like mammalian ATG8s (mATG8s) to phosphatidylethanolamine (PE) to enable mATG8 interaction with the phagophore membrane. The lipid transfer complex includes ATG2, ATG9, and WIPI4/WDR45 and facilitates rapid transfer of lipids to the growing phagophore membrane. When autophagosome biogenesis is complete, the autophagosome biogenesis machinery dissociates from the membrane, leaving the autophagosome labeled with mATG8s. The autophagosome is sealed and later fuses with the endolysosomal system for degradation of autophagic contents.

We previously interrogated how autophagy changes in dorsal root ganglion (DRG) neurons during aging. We discovered that the rate of autophagosome biogenesis decreases drastically with age in primary murine DRG neurons, but that the recruitment of autophagosome biogenesis complexes was not altered with age. Importantly, ectopically expressing WIPI2B in neurons from aged mice restored the rate of autophagosome biogenesis to that of neurons from young adult mice. Further, this restoration depends on the phosphorylation state of WIPI2B at serine 395; local dephosphorylation of WIPI2B S395 at the developing autophagosome is required for autophagosome biogenesis to proceed, independent of age (Stavoe et al., 2019).

We have previously examined autophagy complexes and components across aging (Stavoe et al., 2019; Tsong et al., 2023). However, the potential role played by biological sex in neuronal autophagy was not addressed in those or, to our knowledge, any other studies. Here we re-analyze our previously published data (Stavoe et al., 2019; Tsong et al., 2023) to interrogate how biological sex influences autophagosome biogenesis in murine neurons across development and aging. We examine representative components of the autophagosome biogenesis complexes in multi-color, live-cell imaging of primary DRG neurons from four ages across the murine lifespan, comparing results from female and male mice. We further assay the protein levels of 22 autophagy components by immunoblot of brain and DRG lysates from female and male mice of four life stages. Ultimately, we find sex-related difference in protein levels in only 16% of comparisons, with only modest differences in the comparisons that reached statistical significance. Congruently, we observe no differences in autophagosome formation or recruitment of autophagy components between DRG neurons taken from female and male mice at any age examined. Therefore, our data indicate that the low fraction of sex-related differences in protein levels of autophagy components are not sufficient to produce sex-related differences in autophagosome formation.

## RESULTS

### The autophagy induction complex is not affected by biological sex in neurons

We first examined the induction step of autophagosome biogenesis (Figure 1A) in DRG neurons from male and female mice. We ectopically expressed a labeled component of the induction machinery, mCherry-ATG13, in DRG neurons and quantified the number of mCh-ATG13 puncta at the distal end of DRG neurites taken from male and female mice of three different ages: 3-month-old, young adult mice; 16-17-month-old, aged mice; and 24-month-old, advanced aged mice (Figure 1B-C). We previously found that mCh-ATG13 puncta did not change across ages, independent of sex (Stavoe et al., 2019). Similarly, we detected no differences in mCh-ATG13 puncta between DRG neurons from female and male mice at any of the three ages we examined (Figure 1D), suggesting that sex does not influence the induction step of autophagosome biogenesis in DRG neurons.

**Figure 1.**
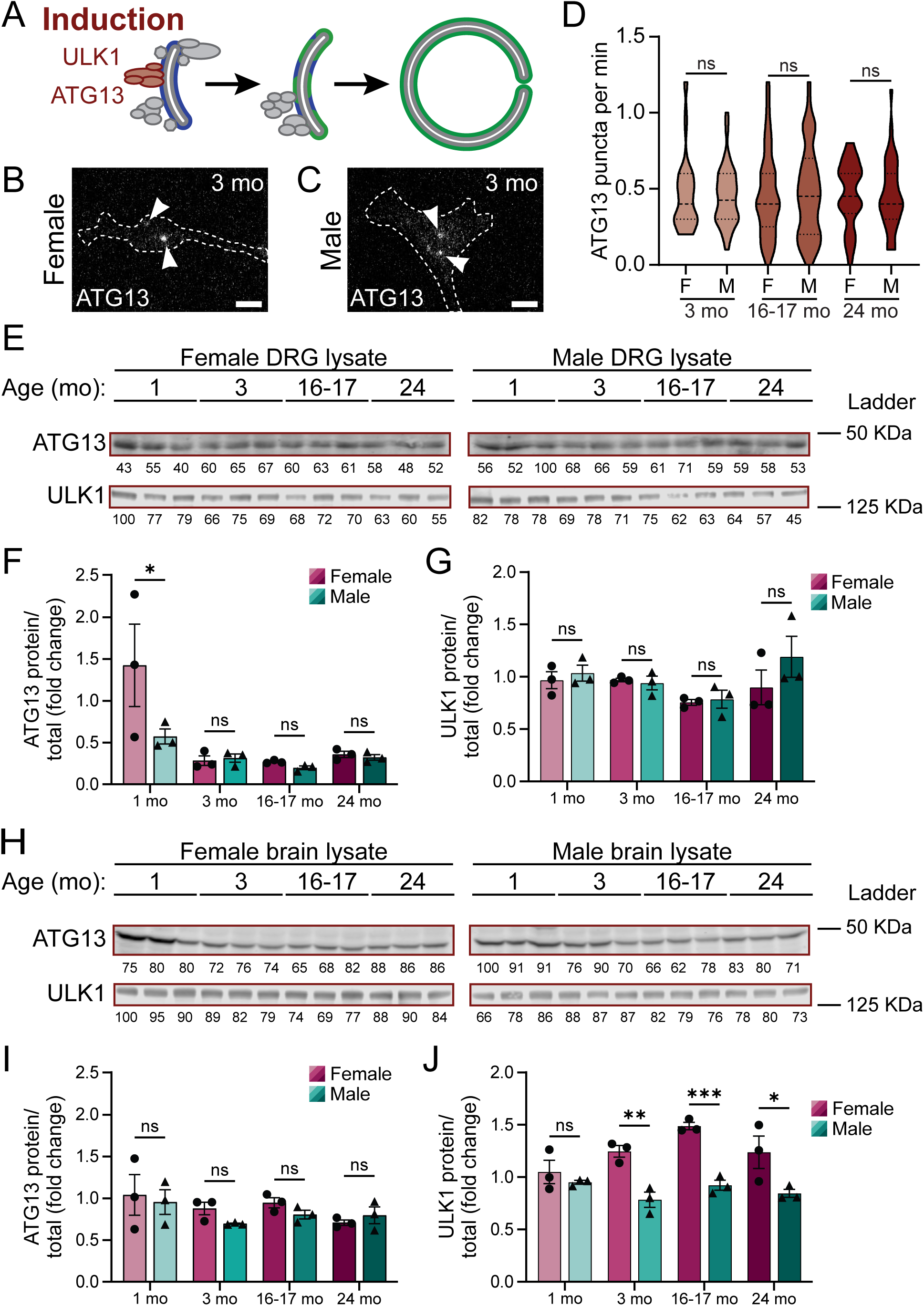
Sex does not affect the induction stage of autophagosome biogenesis. (A) Schematic of autophagosome biogenesis pathway, highlighting the induction complex. (B-C) Representative micrographs of DRG neurons transiently expressing mCh-ATG13 from female (B) and male (C) young adult mice. Arrows indicate mCh-ATG13 puncta. Scale bars, 2 μm. (D) Quantification of the rate of mCh-ATG13 puncta formation in live-cell imaging of DRG neurons (median ± quartiles; n ≥ 34 neurons from three biological replicates of each sex). ns, not significant by unpaired t tests (p > 0.89). (E) Immunoblots of DRG lysates from young (1 mo), young adult (3 mo), aged (16-17 mo), and advanced aged (24 mo) mice of both sexes (n= 3 biological replicates for each sex at each age. Total protein was used as a loading control; normalization factor is indicated below each blot as a percentage. (F-G) Quantification of protein levels of immunoblots in (E), normalized to total protein and represented as a fold change relative to the mean for lysates from young mice (mean ± sem). ns, not significant; *p < 0.05 by two-way ANOVA with Šídák’s multiple comparisons test. (H) Immunoblots of brain lysates from young, young adult, aged, and advanced aged mice of both sexes (n= 3 biological replicates for each sex at each age. Total protein was used as a loading control; normalization factor is indicated below each blot as a percentage. (I-J) Quantification of protein levels of immunoblots in (H), normalized to total protein and represented as a fold change relative to the mean for lysates from young mice (mean ± sem). ns, not significant; *p < 0.05; **p < 0.005; ***p < 0.0005 by two-way ANOVA with Šídák’s multiple comparisons test.

We next asked whether induction complex components had altered protein levels between the sexes in DRG lysates across four different ages: 1-month-old, young mice; 3-month-old, young adult mice; 16-17-month-old, aged mice; and 24-month-old, advanced aged mice. We discerned a difference in ATG13 protein levels in DRG lysates only between female and male young (1 mo) mice; we observed no sex-related differences in ATG13 protein levels in the lysates from adult animals (**Figure *1***E-F). ULK1 is serine/threonine kinase in the induction complex that is itself regulated by phosphorylation. mTOR phosphorylates ULK1 at serine 757 (S757) which inhibits autophagosome biogenesis (Kim et al., 2011). We saw no sex-dependent differences total ULK1 protein (Figure 1E, G) or pULK1(S757) (Figure S1A-B) between DRG lysates harvested from mice at each age.

We also examined whether induction complex components had different protein levels between the sexes in whole brain lysates across the same four ages that were assessed in DRG lysates. We identified no differences in ATG13 protein levels between brain lysates of female and male mice at any age tested (**Figure *1***H-I). We observed that whole brain lysate harvested from female mice had significantly higher levels of total ULK1 in young adult, aged and advanced aged mice, while we did not detect any differences in total ULK1 levels between female and male brain lysates in young mice (Figure 1H, J). Intriguingly, we saw the opposite for pULK1(S757) levels, with no differences between female and male brain lysates in young adult, aged, and advanced aged mice, but males had higher levels of pULK1(S757) in young brain lysate (Figure S1C-D). Given that we identified no sex-related changes in ULK1 or pULK1 levels in DRG lysates, the differences we observed in whole brain lysate might be due to non-neuronal tissues present in the lysates or fundamental differences between the regulation of autophagy protein levels in the central and peripheral nervous systems. Taken together, our results indicated that biological sex does not influence the autophagy induction complex.

### Biological sex does not alter the autophagy nucleation complex in neurons

We next examined the nucleation complex of autophagosome biogenesis, which generates PI3P at the growing phagophore (Figure 2A). To visualize the nucleation complex, we ectopically expressed Halo-ATG14 in isolated DRG neurons (Tsong et al., 2023). We quantified the number of ATG14 puncta in the distal DRG neurites from female and male advanced aged mice (Figure 2B-C). As with the induction complex, we previously found that Halo-ATG14 recruitment did not change across ages, independent of sex (Tsong et al., 2023). Similarly, we saw no sex differences in the rate of Halo-ATG14 puncta in DRG neurons from advanced aged mice (Figure 2D), suggesting that sex does not influence the nucleation step of autophagosome biogenesis in DRG neurons.

**Figure 2.**
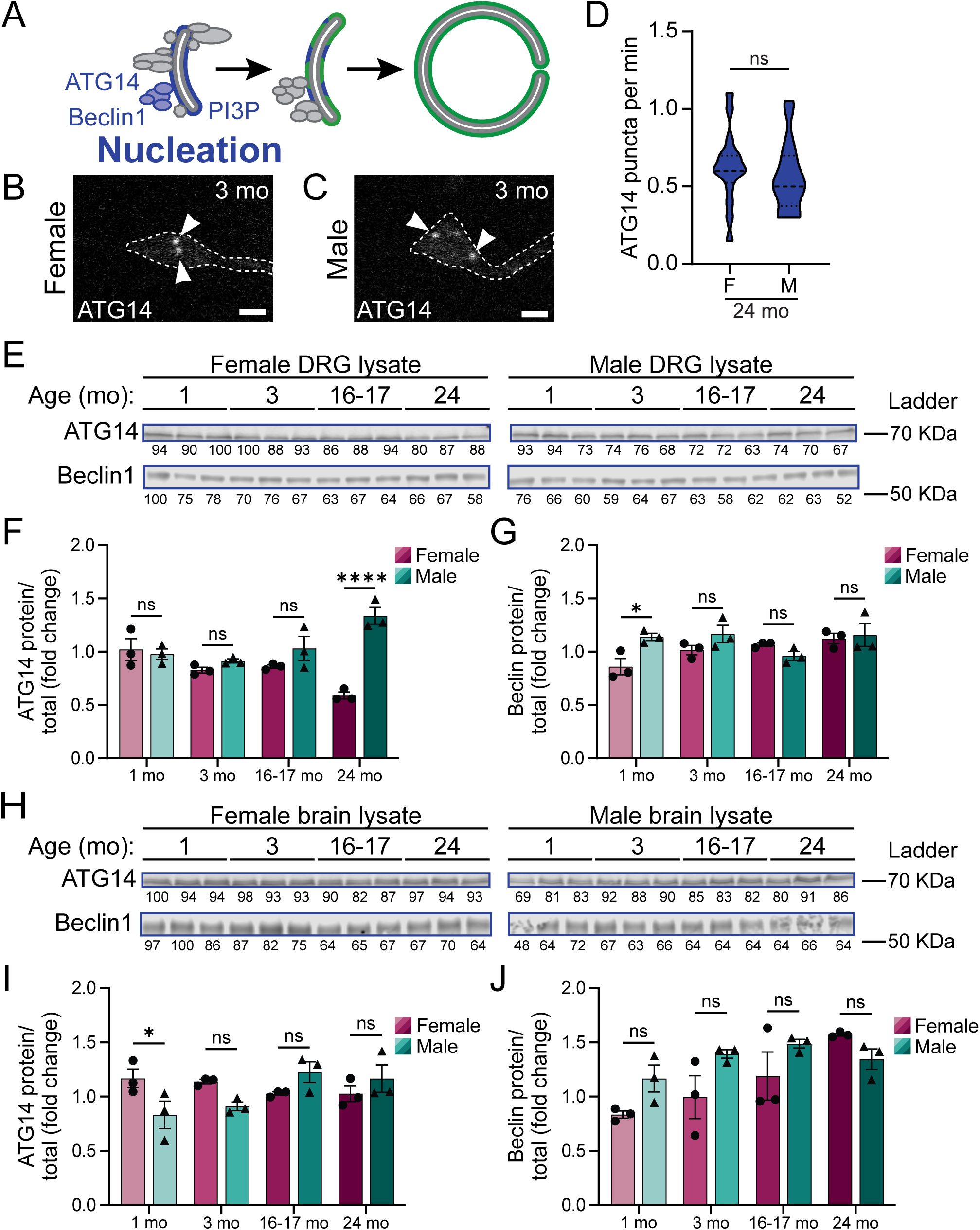
Sex does not affect the nucleation stage of autophagosome biogenesis. (A) Schematic of autophagosome biogenesis pathway, highlighting the nucleation complex. (B-C) Representative micrographs of DRG neurons transiently expressing Halo-ATG14 from female (B) and male (C) young adult mice. Arrows indicate Halo-ATG14 puncta. Scale bars, 2 μm. (D) Quantification of the rate of Halo-ATG14 puncta formation in live-cell imaging of DRG neurons (median ± quartiles; n ≥ 16 neurons from three biological replicates of each sex). ns, not significant by unpaired t tests (p = 0.40). (E) Immunoblots of DRG lysates from young, young adult, aged, and advanced aged mice of both sexes (n= 3 biological replicates for each sex at each age. Total protein was used as a loading control; normalization factor is indicated below each blot as a percentage. (F-G) Quantification of protein levels of immunoblots in (E), normalized to total protein and represented as a fold change relative to the mean for lysates from young mice (mean ± sem). ns, not significant; *p < 0.05; ****p < 0.0001 by two-way ANOVA with Šídák’s multiple comparisons test. (H) Immunoblots of brain lysates from young, young adult, aged, and advanced aged mice of both sexes (n= 3 biological replicates for each sex at each age. Total protein was used as a loading control; normalization factor is indicated below each blot as a percentage. (I-J) Quantification of protein levels of immunoblots in (H), normalized to total protein and represented as a fold change relative to the mean for lysates from young mice (mean ± sem). ns, not significant; *p < 0.05 by two-way ANOVA with Šídák’s multiple comparisons test.

We next interrogated whether nucleation complex components had altered protein levels between female and male mice across development and aging. Upon autophagy induction in mammalian cell culture, ULK1 phosphorylates ATG14 at serine 29 (S29), facilitating autophagosome formation (Park et al., 2016). We examined total ATG14 and pATG14(S29) protein levels in DRG lysates. We observed no differences between female and male DRG lysates from young, young adult, or aged mice. Conversely, in advanced aged mice, total ATG14 protein levels were approximately twice as high in male DRG lysate than in female DRG lysate (Figure 2E-F). We detected no differences in pATG14(S29) protein levels between female and male DRG lysates across all four ages (Figure S2A-B). We also assessed levels of Beclin1, discerning a modest difference between the sexes only in the young DRG lysates: male DRG lysate had higher Beclin1 levels than female DRG lysate. The three adult ages displayed no differences in Beclin1 levels between the sexes (Figure 2E, G). Taken together, our results indicate that despite different levels of ATG14 at the advanced age, the nucleation complex of autophagosome biogenesis is not affected by sex across ages in DRG neurons.

We also examined whether nucleation complex component protein levels were affected by biological sex in whole brain lysates. We assessed total ATG14, pATG14(S29), and Beclin1 levels in brain lysates from female and male mice from the four ages. Broadly, we did not observe sex-specific differences in the protein levels of nucleation complex components in brain lysate across the four ages (Figure 2H-J, Figure S1C-D). We identified higher levels of total ATG14 in female compared to male brain lysate only in brains from young mice (Figure 2H-I). Additionally, pATG14(S29) protein levels were higher in male brain lysate compared to female brain lysate from aged mice (Figure S2C-D).

The product of the nucleation complex is PI3P, a signaling lipid that interacts with many autophagosome biogenesis components. Double FYVE-containing protein 1 (DFCP1) binds to PI3P. We examined DFCP1 recruitment to autophagosomes by ectopically expressing Halo-DFCP1 in DRG neurons. Similar to our ATG14 results, independent of sex, we previously did not observe any age-related changes in DFCP1 recruitment to phagophores in DRG neurons (Stavoe et al., 2019). We uncovered no differences in DFCP1 puncta in distal DRG neurites between female and male aged (16-17 mo) mice (Figure S2E). Taken together, our results indicate that the function of the nucleation complex appears unaffected by biological sex in DRG neurons.

### The autophagy elongation complex is not influenced by biological sex in neurons

The elongation complex of autophagosome biogenesis is composed of two ubiquitin-like conjugation complexes which ultimately conjugate phosphatidylethanolamine (PE) to LC3 to enable LC3-II interaction with the phagophore (Tanida et al., 2004) (Figure 3A). To visualize the elongation complex, we ectopically expressed mCh-ATG5 in DRG neurons harvested from male and female mice (Figure 3B-C). We quantified mCh-ATG5 puncta in DRG distal neurites from female and male young adult, aged, and advanced aged mice. Similar to our findings for the induction and nucleation complexes, we did not observe any differences in ATG5 recruitment between DRG neurons from female and male mice across the three ages (Figure 3D).

**Figure 3.**
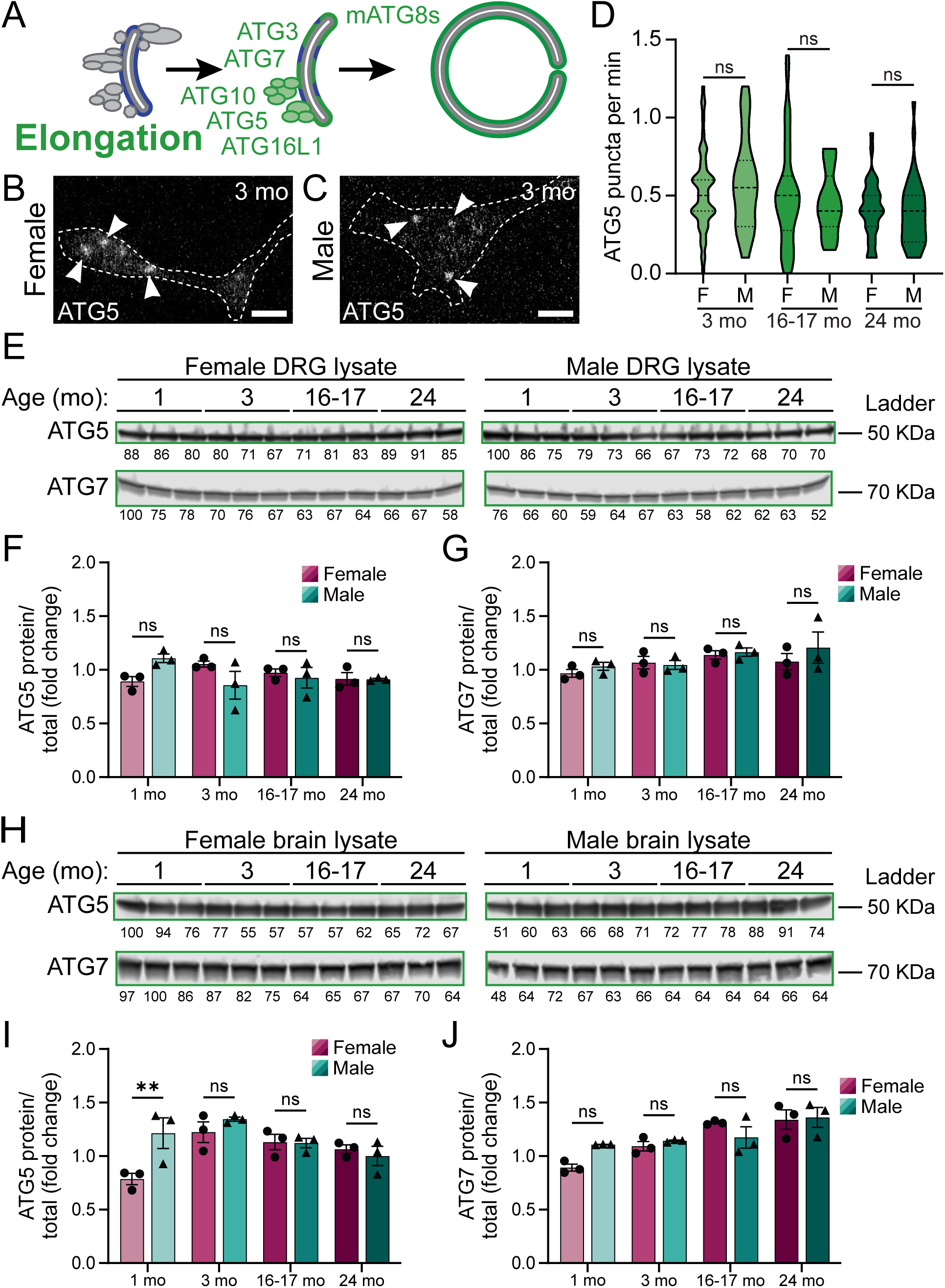
Sex does not affect the elongation machinery of autophagosome biogenesis. (A) Schematic of autophagosome biogenesis pathway, highlighting the elongation complex. (B-C) Representative micrographs of DRG neurons transiently expressing mCh-ATG5 from female (B) and male (C) young adult mice. Arrows indicate mCh-ATG5 puncta. Scale bars, 2 μm. (D) Quantification of the rate of mCh-ATG5 puncta formation in live-cell imaging of DRG neurons (median ± quartiles; n ≥ 22 neurons from three biological replicates of each sex). ns, not significant by unpaired t tests (p > 0.39). (E) Immunoblots of DRG lysates from young, young adult, aged, and advanced aged mice of both sexes (n= 3 biological replicates for each sex at each age. Total protein was used as a loading control; normalization factor is indicated below each blot as a percentage. (F-G) Quantification of protein levels of immunoblots in (E), normalized to total protein and represented as a fold change relative to the mean for lysates from young mice (mean ± sem). ns, not significant by two-way ANOVA with Šídák’s multiple comparisons test. (H) Immunoblots of brain lysates from young, young adult, aged, and advanced aged mice of both sexes (n= 3 biological replicates for each sex at each age. Total protein was used as a loading control; normalization factor is indicated below each blot as a percentage. (I-J) Quantification of protein levels of immunoblots in (H), normalized to total protein and represented as a fold change relative to the mean for lysates from young mice (mean ± sem). ns, not significant; **p < 0.005 by two-way ANOVA with Šídák’s multiple comparisons test.

We next asked whether elongation complex components had altered protein levels between female and male mice across development and aging. We examined ATG3, ATG5, ATG7, ATG10, and ATG16L1 protein levels in DRG lysates (Figure 3E-G; Figure S3A-D). We observed no differences in ATG5, ATG7, or ATG16L1 protein levels between female and male DRG lysates at any age (Figure 3E-G; Figure S3A, D). For ATG3 protein levels, we only detected a difference between females and males in young DRG lysates (Figure S3A-B). Additionally, for ATG10 protein levels, we found statistically significant differences between females and males only in advanced aged DRG lysates (Figure S3A, C). Thus, elongation complex components do not exhibit gross differences in protein levels between female and male DRG lysates.

We also examined whether elongation complex component protein levels were affected by biological sex in whole brain lysates (Figure 3H-J; Figure S3E-H). For ATG5 protein levels, we detected a sex-specific difference only in young brain lysate, with higher ATG5 protein levels in male compared to female brain lysate from young mice (Figure 3H-I). We discerned no differences in ATG7 protein levels between female and male brain lysate at any age (Figure 3H, J). In addition, we observed that male brain lysate had modestly higher ATG3 levels than female brain lysate in young and young adult lysates, but no differences in aged or advanced aged lysates (Figure S3E-F). Likewise, we saw a difference in ATG10 protein levels between female and male lysates only in samples from young adult mice (Figure S3E, G). We also discovered modestly higher ATG16L1 levels in male compared to female brain lysate from young mice, but no differences in any of the adult lysates (Figure S3E, H). Taken together, our results suggest that the elongation complex is not different between the sexes in neurons across development and aging.

### Biological sex does not influence autophagosome formation in neurons across development and aging

The ultimate function of the elongation complex is to conjugate PE to LC3B and its homologs, collectively called mammalian ATG8s (mATG8s) (Figure 4A). We previously discovered a profound decrease in the formation of autophagic vesicles (AVs) in DRG neurons with age, independent of sex (Stavoe et al., 2019). We used the GFP-LC3B probe to examine the rate of autophagosome biogenesis across four ages of female and male mice (Figure 4B-C). We quantified the rate of AV biogenesis events by tracking the formation of discrete GFP-LC3B puncta. We observed no differences in the rate of AV formation between DRG neurons from female and male mice at each age examined (Figure 4D-E), in keeping with our data from the induction, nucleation, and elongation complexes.

**Figure 4.**
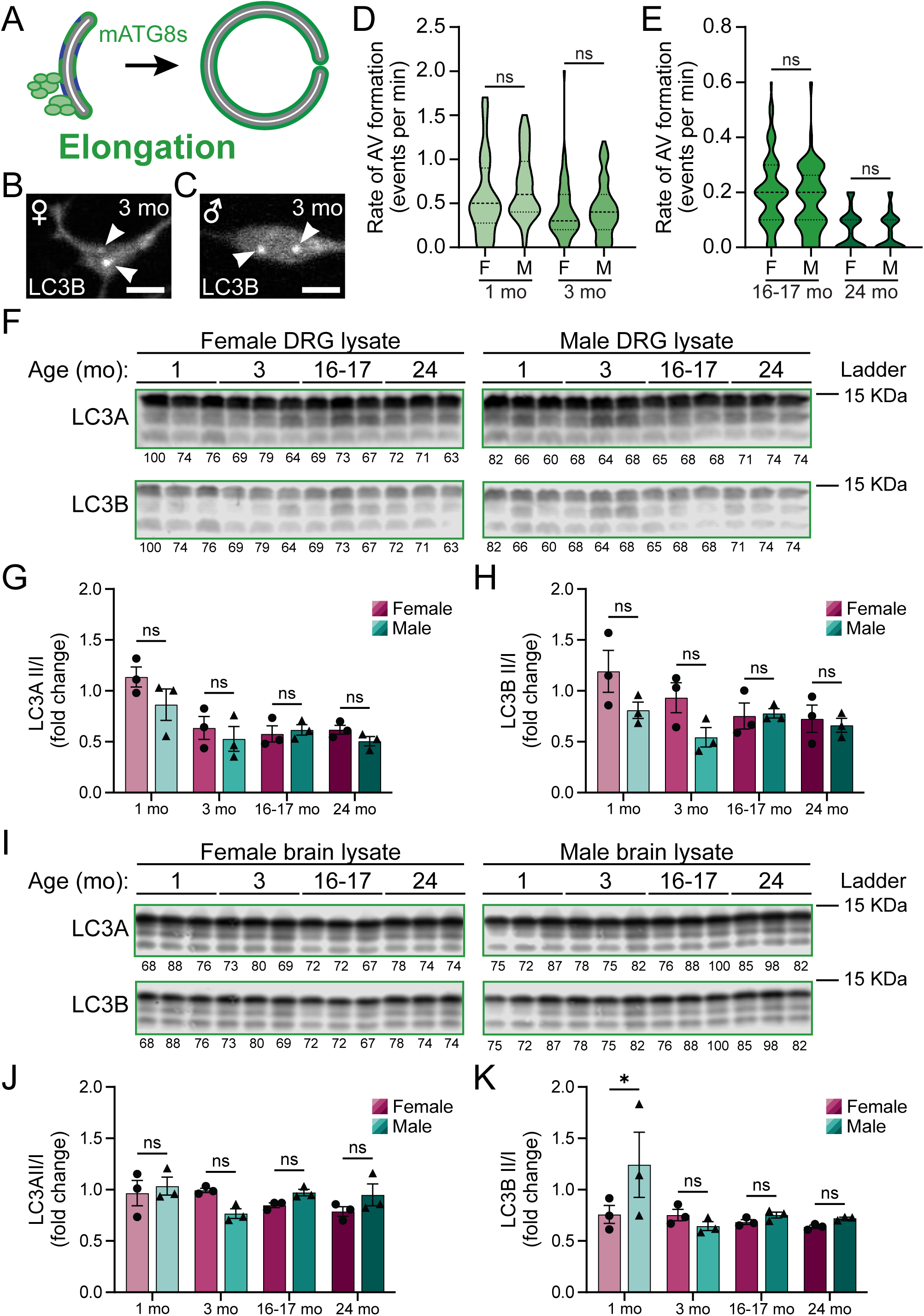
Sex does not affect mATG8s nor autophagosome biogenesis. (A) Schematic of autophagosome biogenesis pathway, highlighting the elongation complex. (B-C) Representative micrographs of DRG neurons expressing GFP-LC3B from female (B) and male (C) young adult mice. Arrows indicate GFP-LC3B puncta (autophagic vesicles). Scale bars, 2 μm. (D-E) Quantification of the rate of autophagic vesicle formation in live-cell imaging of DRG neurons (median ± quartiles; n ≥ 30 neurons from three biological replicates of each sex). ns, not significant by unpaired t tests (p > 0.12). (F) Immunoblots of DRG lysates from young, young adult, aged, and advanced aged mice of both sexes (n= 3 biological replicates for each sex at each age. Total protein was used as a loading control; normalization factor is indicated below each blot as a percentage. (G-H) Quantification of protein levels of immunoblots in (F), normalized to total protein and represented as a fold change relative to the mean for lysates from young mice (mean ± sem). ns, not significant by two-way ANOVA with Šídák’s multiple comparisons test. (I) Immunoblots of brain lysates from young, young adult, aged, and advanced aged mice of both sexes (n= 3 biological replicates for each sex at each age. Total protein was used as a loading control; normalization factor is indicated below each blot as a percentage. (J-K) Quantification of protein levels of immunoblots in (I), normalized to total protein and represented as a fold change relative to the mean for lysates from young mice (mean ± sem). ns, not significant; *p < 0.05 by two-way ANOVA with Šídák’s multiple comparisons test.

We next asked whether mATG8s had altered protein levels between female and male mice across development and aging. The elongation complex processes mATG8s from pro-mATG8s through to their conjugation to PE, resulting in multiple mATG8 species on an immunoblot. The ratio of the mATG8-II species to the mATG8-I species provides an approximate readout for autophagic activity. Mice express four orthologs of LC3B: γ-aminobutyric acid receptor-associated protein (GABARAP), GABARAP-Like 1 (GABARAPL1/GEC1), and GABARAP-Like 2 (GABARAPL2/GATE16) (Schaaf et al., 2016). Mice do not appear to express LC3C (Liu et al., 2017). In DRG lysates, we detected no differences in LC3A II/I or LC3B II/I ratios between samples from female and male mice across all four ages tested (Figure 4F-H). Similarly, we also observed minimal differences in GABARAP, GABARAPL1, and GABARAPL2 levels between female and male DRG lysates (Figure S4). We saw a slight difference in GABARAP levels between female and male samples only in young DRG lysates (Figure S4A-B). Congruently, we discerned a significant difference in the GABARAPL1 II/I ratio between female and male samples only from aged mice (Figure S4A, C). We uncovered no differences in GABARAPL2 levels between female and male DRG lysates from all ages examined (Figure S4A, D).

We further assayed whether mATG8 protein levels were affected by sex in whole brain lysates (Figure 4I-K; Figure S5). We observed no differences in LC3A (Figure 4I-H), GABARAP (Figure S5A-B), GABARAPL1 (Figure S5A, C), or GAABRAPL2 (Figure S5A, D) protein levels between female and male brain lysate at any age. We identified a difference in the LC3B II/I ratio between female and male samples only from young brain lysates (Figure 4I, K).

The autophagy adapter protein p62 delivers cargo to an autophagosome by interacting with aggregated proteins and mATG8s at the growing phagophore (Pankiv et al., 2007). It is degraded along with the cargo once the autophagosome fuses with the endolysosomal system; thus, p62 is used as a readout for autophagic activity (Mizushima et al., 2010). While we observed differences in p62 levels between female and male DRG lysates from young, young adult, and aged mice (Figure S4A, E), we detected a sex-based difference in p62 levels only in brain lysates from advanced aged mice (Figure S5A, E). Taken together, our data indicate that autophagosome formation is not altered in neurons between females and males across development and aging.

### WIPI family members are not different in neurons from male and female mice

The WD-repeat protein interacting with phosphoinositides (WIPI) family has four family members in mammals, WIPI1-WIPI4 (WIPI3 and WIPI4 are also known as WDR45B and WDR45, respectively) (Polson et al., 2010; Proikas-Cezanne et al., 2004). WIPIs play essential roles in autophagosome formation, bridging PI3P on the phagophore membrane with the autophagosome biogenesis complexes (Figure 5A) (Dooley et al., 2014). We previously showed that ectopic expression of WIPI2 can restore the rate of autophagosome biogenesis in DRG neurons from aged mice to the rate observed in DRG neurons from young adult mice (Stavoe et al., 2019), so we were interested to interrogate whether biological sex influences WIPI proteins. We first transiently expressed SNAP-WIPI1 in DRG neurons from female and male aged mice (Figure 5B-C). We observed no difference in the rate of WIPI1 recruitment between DRG neurons from female and male aged mice (Figure 5D).

**Figure 5.**
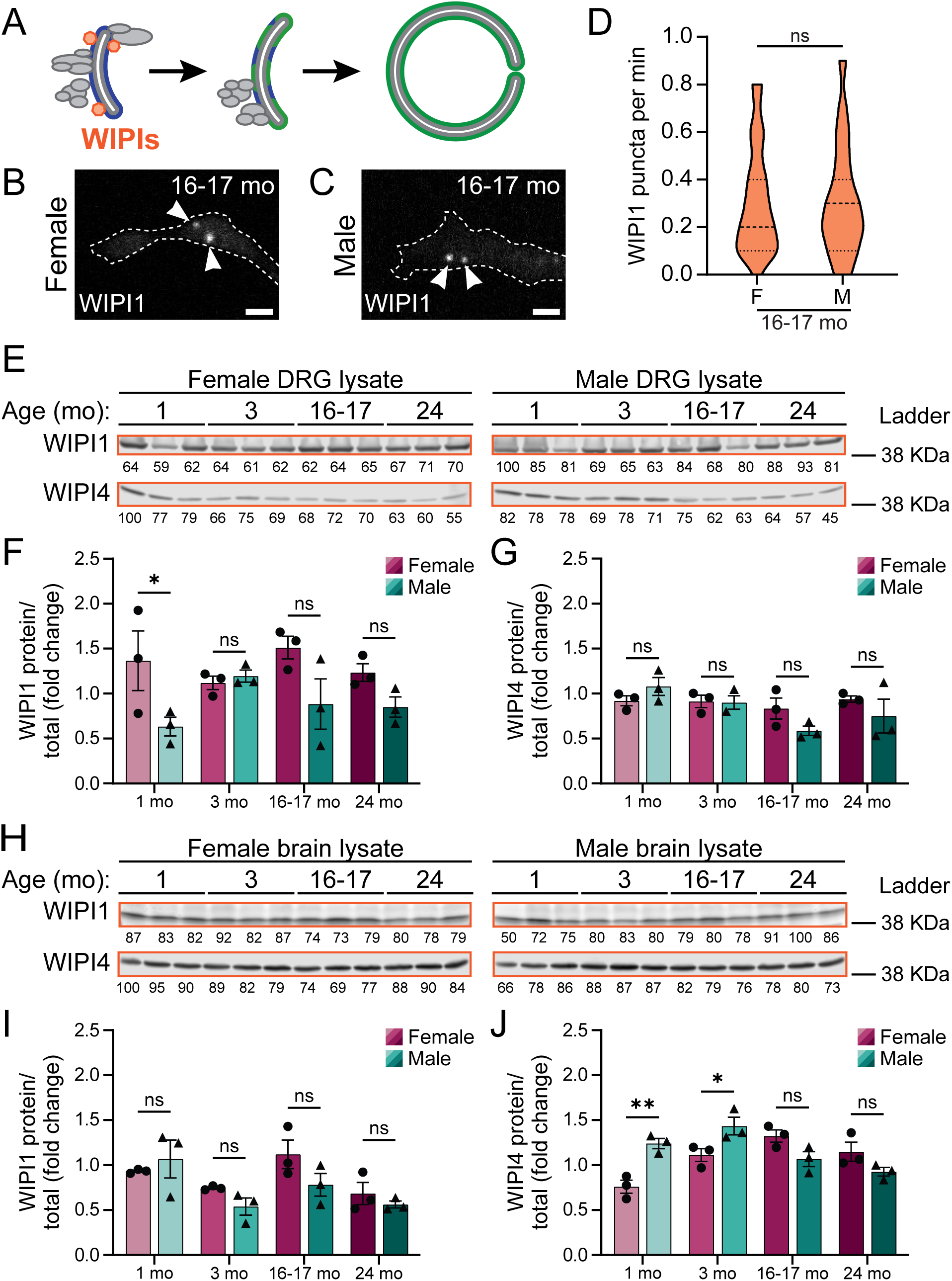
Sex does not affect WIPIs. (A) Schematic of autophagosome biogenesis pathway, highlighting WIPI proteins. (B-C) Representative micrographs of DRG neurons transiently expressing SNAP-WIPI1 from female (B) and male (C) aged mice. Arrows indicate SNAP-WIPI1 puncta. Scale bars, 2 μm. (D) Quantification of the rate of SNAP-WIPI1 puncta formation in live-cell imaging of DRG neurons (median ± quartiles; n = 43 neurons from three biological replicates of each sex). ns, not significant by unpaired t tests (p = 0.96). (E) Immunoblots of DRG lysates from young, young adult, aged, and advanced aged mice of both sexes (n= 3 biological replicates for each sex at each age. Total protein was used as a loading control; normalization factor is indicated below each blot as a percentage. (F-G) Quantification of protein levels of immunoblots in (E), normalized to total protein and represented as a fold change relative to the mean for lysates from young mice (mean ± sem). ns, not significant; *p < 0.05 by two-way ANOVA with Šídák’s multiple comparisons test. (H) Immunoblots of brain lysates from young, young adult, aged, and advanced aged mice of both sexes (n= 3 biological replicates for each sex at each age. Total protein was used as a loading control; normalization factor is indicated below each blot as a percentage. (I-J) Quantification of protein levels of immunoblots in (H), normalized to total protein and represented as a fold change relative to the mean for lysates from young mice (mean ± sem). ns, not significant; *p < 0.05; **p < 0.005 by two-way ANOVA with Šídák’s multiple comparisons test.

We next asked whether WIPIs had altered protein levels between female and male mice across development and aging. In DRG lysates, we detected a sex-related difference in WIPI1 protein levels only between lysates from female and male young mice; we observed no sex-related differences in WIPI1 protein levels in the lysates from adult animals (Figure 5E-F). Additionally, we saw differences in WIPI3 protein levels between female and male DRG lysates from young and young adult mice, but not from aged or advanced aged mice (Figure S6A-B). We identified no differences in WIPI4 protein levels between female and male lysates from mice of any age assayed (Figure 5E, G).

We also examined whether WIPI protein levels were affected by sex in whole brain lysates. We observed no differences in WIPI1 (Figure 5H-I) or WIPI3 (Figure S6C-D) protein levels between female and male brain lysates at any age. Conversely, we discerned differences in WIPI4 protein levels between female and male brain lysates from young and young adult mice, but not from aged or advanced aged mice (Figure 5H, J).

Given the unique ability of WIPI2B to restore age-induced autophagosome biogenesis decline in DRG neurons (Stavoe et al., 2019), we closely examined whether WIPI2 was affected by biological sex across development and aging. We first ectopically expressed Halo-WIPI2B in DRG neurons from aged animals to assess WIPI2 recruitment (Figure 6A-C). We detected no differences in the rate of Halo-WIPI2B recruitment between DRG neurons from female and male aged mice (Figure 6D). In addition, we assessed how biological sex influenced restoration of AV formation by ectopic expression of WIPI2B in neurons from aged mice with the GFP-LC3B autophagosome probe. In neurons from aged mice, biological sex did not affect the rate of AV formation, consistent with our earlier data (**Figure *4***C). Congruently, in neurons from aged mice with ectopic expression of WIPI2B, biological sex did not affect the restoration of AV formation, with neurons from both female and male mice displaying higher rates of autophagosome biogenesis than control neurons. Importantly, the rate of AV formation in neurons ectopically expressing WIPI2B was not different between neurons from female and male aged mice (**Figure *6***E).

**Figure 6.**
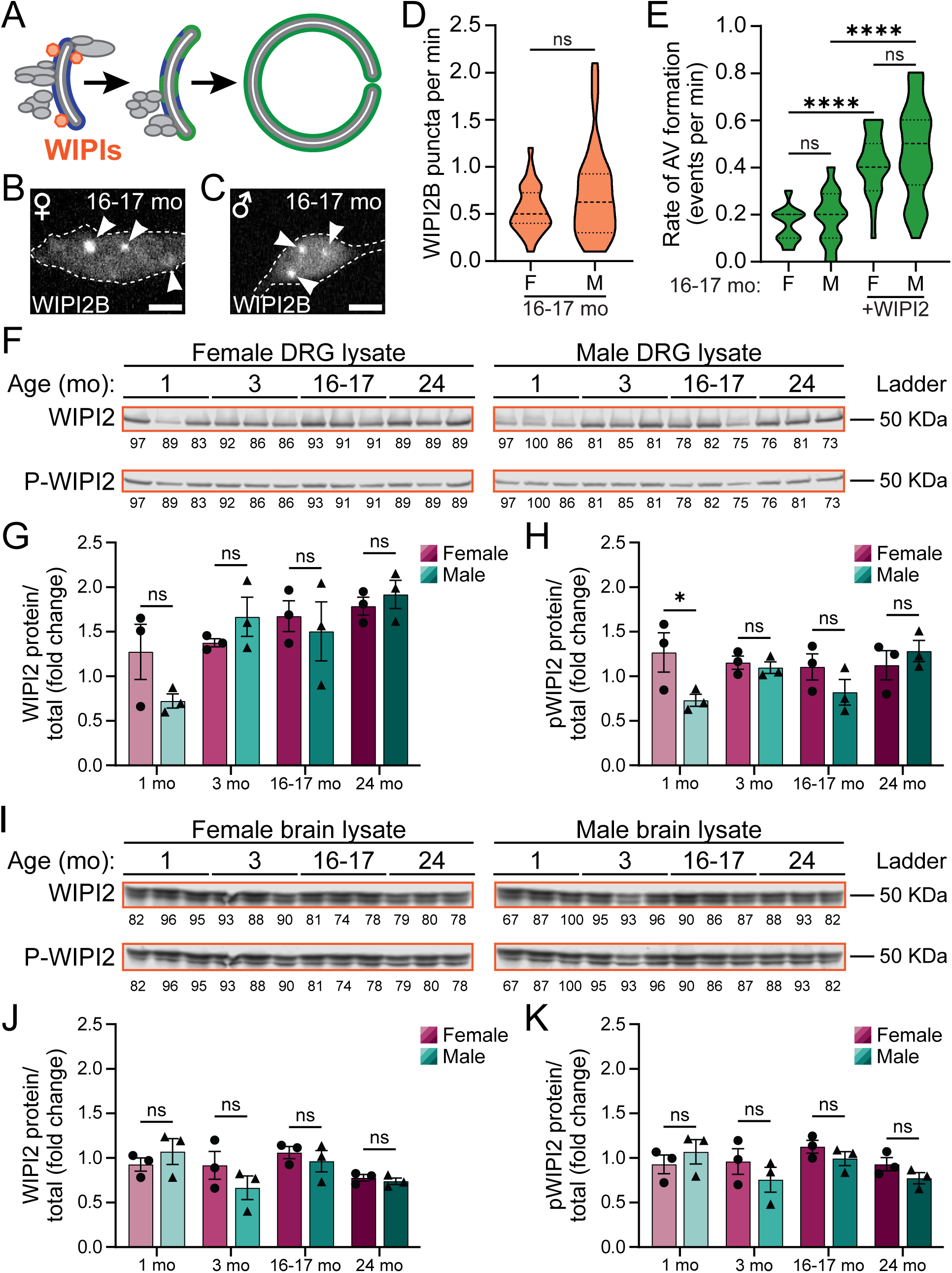
Sex does not affect WIPI2 recruitment nor its ability to restore autophagosome biogenesis in neurons from aged mice. (A) Schematic of autophagosome biogenesis pathway, highlighting WIPI proteins. (B-C) Representative micrographs of DRG neurons transiently expressing Halo-WIPI2B from female (B) and male (C) aged mice. Arrows indicate Halo-WIPI2B puncta. Scale bars, 2 μm. (D) Quantification of the rate of Halo-WIPI2B puncta formation in live-cell imaging of DRG neurons (median ± quartiles; n = 30 neurons from ≥ 5 biological replicates of each sex). ns, not significant by unpaired t tests (p = 0.07). (E) Quantification of the rate of autophagic vesicle formation in live-cell imaging of DRG neurons (median ± quartiles; n ≥ 23 neurons from ≥ 4 biological replicates of each sex). ns, not significant; ****p < 0.0001 by two-way ANOVA with Tukey’s multiple comparisons test. (F) Immunoblots of DRG lysates from young, young adult, aged, and advanced aged mice of both sexes (n= 3 biological replicates for each sex at each age. Total protein was used as a loading control; normalization factor is indicated below each blot as a percentage. (G-H) Quantification of protein levels of immunoblots in (F), normalized to total protein and represented as a fold change relative to the mean for lysates from young mice (mean ± sem). ns, not significant; *p < 0.05 by two-way ANOVA with Šídák’s multiple comparisons test. (I) Immunoblots of brain lysates from young, young adult, aged, and advanced aged mice of both sexes (n= 3 biological replicates for each sex at each age. Total protein was used as a loading control; normalization factor is indicated below each blot as a percentage. (J-K) Quantification of protein levels of immunoblots in (I), normalized to total protein and represented as a fold change relative to the mean for lysates from young mice (mean ± sem). ns, not significant by two-way ANOVA with Šídák’s multiple comparisons test.

We further assessed whether WIPI2 protein levels were affected by biological sex in DRG (Figure 6F-H) and whole brain (Figure 6I-K) lysates. WIPI2B is phosphorylated at S395; this phosphorylation regulates the ability of WIPI2B to restore autophagosome biogenesis in DRG neurons from aged mice (Stavoe et al., 2019). We discerned no differences in total WIPI2 protein levels between female and male DRG lysates from all four ages of mice (Figure 6F-G). We saw a difference in pWIPI2 (S395) levels between female and male DRG lysates only at the young age (Figure 6F, H). Similarly, we observed no differences in total WIPI2 or pWIPI2 (S395) (Figure 6I-K) protein levels between female and male brain lysates at any age. Together, these data indicate that biological sex does not influence recruitment of WIPI proteins to the phagophore in neurons, nor the ability of WIPI2B to restore age-related declines in neuronal autophagosome formation.

## DISCUSSION

Here, we interrogated whether biological sex affected autophagosome formation in murine neurons during development and aging. We re-analyzed our previous data (Stavoe et al., 2019; Tsong et al., 2023) to discern any differences in autophagy between neurons and tissues harvested from female and male mice. We systematically examined different stages of autophagosome biogenesis and directly assessed the rate of autophagosome formation with multi-color, live-cell imaging of DRG neurons isolated from four ages of female and male mice: 1-month-old, young mice; 3-month-old, young adult mice; 16-17-month-old, aged mice; and 24-month-old, advanced aged mice. We found minimal differences in protein levels of autophagosome biogenesis components in both DRG and brain lysates between female and male mice of all four ages tested. Importantly, we did not discern any sex-related differences in recruitment of autophagosome biogenesis components ATG13, ATG14, ATG5, WIPI1, or WIPI2B. Ultimately, we did not observe any differences in the rate of autophagosome formation between DRG neurons taken from female or male mice of any of the four ages examined. Thus, we propose that biological sex is not a contributing variable to neuronal autophagosome biogenesis during development or aging.

We analyzed the protein levels of 22 autophagy-related components at four different ages (1 month, 3 months, 16-17 months, and 24 months) in two tissues (DRG and whole brain) from female and male mice, resulting in 176 comparisons between the sexes. We found differences in only 29 (16%) comparisons between female and male tissues: 14 in DRG lysates (8%) and 15 in brain lysates (8%). Across both lysates and all ages, of the 29 comparisons that did display sex-related differences, 55% of the differences had higher protein levels in male compared to female lysates. Interestingly, we discovered that the majority of the sex-related differences were in lysates from young mice (52%), with the remaining differences split roughly evenly across the adult ages (21% at 3 months, 14% at 16-17 months, and 14% at 24 months). Importantly, the 1-month timepoint is well in advance of puberty and accompanying increases in release of gonadal hormones. In addition to hormonal sex differences, brain development is influenced by developmental differences in gene expression between the sexes (Wolstenholme et al., 2013), which may contribute to the larger sex-related variance we detected in autophagy protein levels in lysates from young mice. Importantly, several neurodevelopmental processes are still ongoing in 1-month-old mouse brains, including myelination (Hammelrath et al., 2016), cortical maturation, and synaptic plasticity.

Despite the few differences we identified between the sexes in DRG and brain lysate, we did not detect any sex-related differences in the recruitment of autophagy complexes in DRG neurons. Induction complex component ATG13 (**Figure *1***D) and elongation complex component ATG5 (**Figure *3***D) were recruited at similar rates in DRG neurons between female and male adult mice (3 months, 16-17 months, and 24 months). Similarly, we found no differences in the rate of recruitment of the nucleation complex component ATG14 in DRG neurons between female and male advanced aged mice (**Figure *2***D). Our results were congruent with WIPI1 and WIPI2B in DRG neurons from aged mice, with no differences observed between the sexes (**Figure *5***D, **Figure *6***D). These complexes and proteins function in concert to form autophagosomes, so the rate of autophagosome formation depends on the rate-limiting component. Thus, disparities between the sexes in protein levels of non-rate-limiting components would not likely alter the rate of autophagosome biogenesis.

Ultimately, we did not discover any sex-related differences in the rate of autophagosome formation in DRG neurons from young, young adult, aged, or advanced aged mice (**Figure *4***D-E). Importantly, the ability of ectopic expressed WIPI2B to restore the rate of autophagosome formation in DRG neurons from aged mice similarly did not display sex-related differences (**Figure *6***E). Altogether, our data indicate that biological sex is not a variable that influences autophagosome formation in DRG neurons.

## MATERIALS AND METHODS

The data presented here were predominantly previously collected and published in (Stavoe et al., 2019; Tsong et al., 2023). Here, we reanalyzed that data to assess sex-related differences. New data for this paper include: ATG13 expression levels in DRG and brain lysate; WIPI1 puncta formation; and WIPI2B puncta formation. To achieve appropriate n values for each sex, we collected new data for ATG14 puncta at 24 mo.

### Reagents

GFP-LC3B transgenic mice (strain: B6.Cg-Tg[CAG-EGFP/LC3]53Nmi/NmiRbrc) were generated by N. Mizushima (Tokyo Medical and Dental University, Tokyo, Japan; [29]) and obtained from RIKEN BioResource Center in Japan (RBRC00806). These mice were bred with C57BL/6J mice obtained from The Jackson Laboratory (000664). Hemizygous and wild-type littermates were used in experiments. Constructs used include: mCherry-ATG13 (subcloned from Addgene, 22875; deposited by Noboru Mizushima), mCherry-ATG5 (Addgene 13095), Halo-ATG14L (subcloned from Addgene, 21635; deposited by Tamotsu Yoshimori), SNAP-WIPI1A (subcloned from Addgene 38272), Halo-DFCP1 (subcloned from Addgene 38269), and Halo-WIPI2B (subcloned from GFP-WIPI2B (Dooley et al., 2014)), all previously published in (Stavoe et al., 2019; Tsong et al., 2023).

### Primary neuron culture

Mice were euthanized prior to dissection. All animal protocols were approved by the Institutional Animal Care and Use Committee at the University of Pennsylvania. DRG neurons were isolated as previously described [59] from mice of either sex in these postnatal ranges: P21-28 (1 mo, young), P90-120 (3 mo, young adult), P480-540 (16-17 mo, aged), or P730-761 (24 mo, advanced aged). DRG neurons were plated on glass-bottomed dishes (MatTek Corporation, P35G-1.5-14-C) and cultured in F-12 Ham’s media (Gibco, 11765-047) with 10% heat-inactivated fetal bovine serum (HyCLone, SH30071.03), 100 U/mL penicillin, and 100 μg/mL streptomycin (Gibco, 15140122). DRG neurons were imaged or fixed after being maintained for 2 days at 37°C in a 5% CO_2_ incubator.

Prior to plating, neurons were transfected with a maximum of 0.6 μg total plasmid DNA using a Nucleofector (Lonza, Basel, Switzerland) and following the manufacturer’s instructions. For Halo-ATG14-, Halo-DFCP1-, Halo-WIPI1B-, and SNAP-WIPI1-transfected neurons, DRG neurons were incubated with 100 nM of JF646-Halo ligand (from Luke Levis, Janelia Farms, HHMI) or 100 nM of SNAP-Cell® 647-SiR (New England BioLabs) for at least 30 min at 37°C in a 5% CO_2_ incubator. After incubation, neurons were washed three times with complete equilibrated F-12 media, with the final wash remaining on the neurons for at least 15 min at 37°C in a 5% CO_2_ incubator.

### Live-cell imaging and image analysis

Microscopy was performed in Hibernate A low fluorescence nutrient media (BrainBits, HALF500) supplemented with 2% B27 (Gibco, A3582801) and 2 mM GlutaMAX (Gibco, 35050061). Confocal images were captured with a spinning-disk confocal (UltraVIEW VoX; PerkinElmer, Waltham, MA) microscope (Eclipse Ti; Nikon, Tokyo, Japan) with an Apochromat 100x, 1.49 NA oil immersion objective (Nikon, Tokyo, Japan) at 37°C in an environmental chamber. Digital micrographs were acquired with an EM charge-coupled device camera (C9100; Hammamatsu Photonics, Japan) using Volocity software (PerkinElmer, Waltham, MA). Confocal images for new data (Halo-ATG14 and Halo-DFCP1 at 24 mo) were captured with a spinning-disk confocal (W1 confocal system; Nikon Instruments, Tokyo, Japan) with an Apochromat Lambda 100x, 1.45 NA oil immersion objective (Nikon Instruments, Tokyo, Japan) at 37°C in an environmental chamber. Digital micrographs were acquired with a back-illuminated sCMOS camera (Teledyne Photometrics, Tucson, AZ) using Nikon Elements software (Nikon Instruments, Tokyo, Japan). The Perfect Focus System was used to maintain Z position during time-lapse acquisition on both confocal systems.

To capture autophagosome biogenesis, time-lapse videos were acquired for 10 min with a frame every 3 s. Multiple channels were acquired consecutively, with the green (488 nm) channel captured first, followed by red (561 nm), and far-red (640 nm). DRG neurons were selected for imaging based on morphological criteria and low expression of transfected constructs. To minimize artifacts from overexpression, neurons within a narrow range of low fluorescence intensity were chosen for imaging, ensuring the analyzed neurons expressed low levels of the ectopic tagged proteins.

All image analysis was performed on raw data. Images were prepared in FIJI (Schindelin et al., 2012); contrast and brightness were adjusted equally to all images within a series. Time-lapse micrographs were analyzed with FIJI (Schindelin et al., 2012). To quantify AV biogenesis, GFP-LC3B puncta were tracked manually using FIJI. An AV biogenesis event was defined as the de novo appearance of a GFP-LC3B punctum based on changes in fluorescence intensity over time. For GFP-LC3B puncta that were present at the start of the time-lapse series, only those puncta that increased in fluorescence intensity and/or area with time were counted as AV biogenesis events. Z-stack micrographs were initially processed in FIJI into maximal projections. Maximal projection micrographs were segmented, processed, and subsequently analyzed in FIJI using Analyze Particles.

### Immunoblotting

Brains of non-transgenic mice were dissected and subsequently homogenized and lysed. Brains were homogenized individually in RIPA buffer [1x PBS (see above), 1% Triton X-100 (Fisher, BP151500), 0.5% deoxycholate (Fisher, BP349100), 0.1% SDS (Fisher, BP166500), 1x cOmplete^TM^ protease inhibitor cocktail (Roche, 11836170001), and 1x Halt^TM^ protease and phosphatase inhibitor cocktail (ThermoFisher Scientific, 78440)]. Total protein in each lysate was determined by BCA assay (ThermoFisher Scientific, 23225) to ensure equal protein loading during western blot analysis.

All supernatants were analyzed by SDS-PAGE western blot, transferred onto FL PVDF membranes (Millipore-Sigma, IPFL00010), and visualized with fluorescent secondary antibodies (Li-Cor, ThermoFisher) using an Odyssey® CLx imaging system (Li-Cor, Lincoln, Nebraska). See Key Resources Table for antibodies used. All western blots were analyzed with Image Studio (Li-Cor). Total protein was used as a loading control (REVERT^TM^ Total Protein Stain, Li-Cor, 926-11021). The normalization factor is listed below each blot as a percent.

### Normalization and Statistical Analysis

After normalization to total protein loading, immunoblot data were normalized to the combined (female and male) 1-mo data for each protein assayed. Female and male means at each age were normalized and plotted. Prism 10 (GraphPad) was used to plot graphs and perform statistical tests. Specific statistical tests used are indicated in the text and figure legends.

**Table 1.**
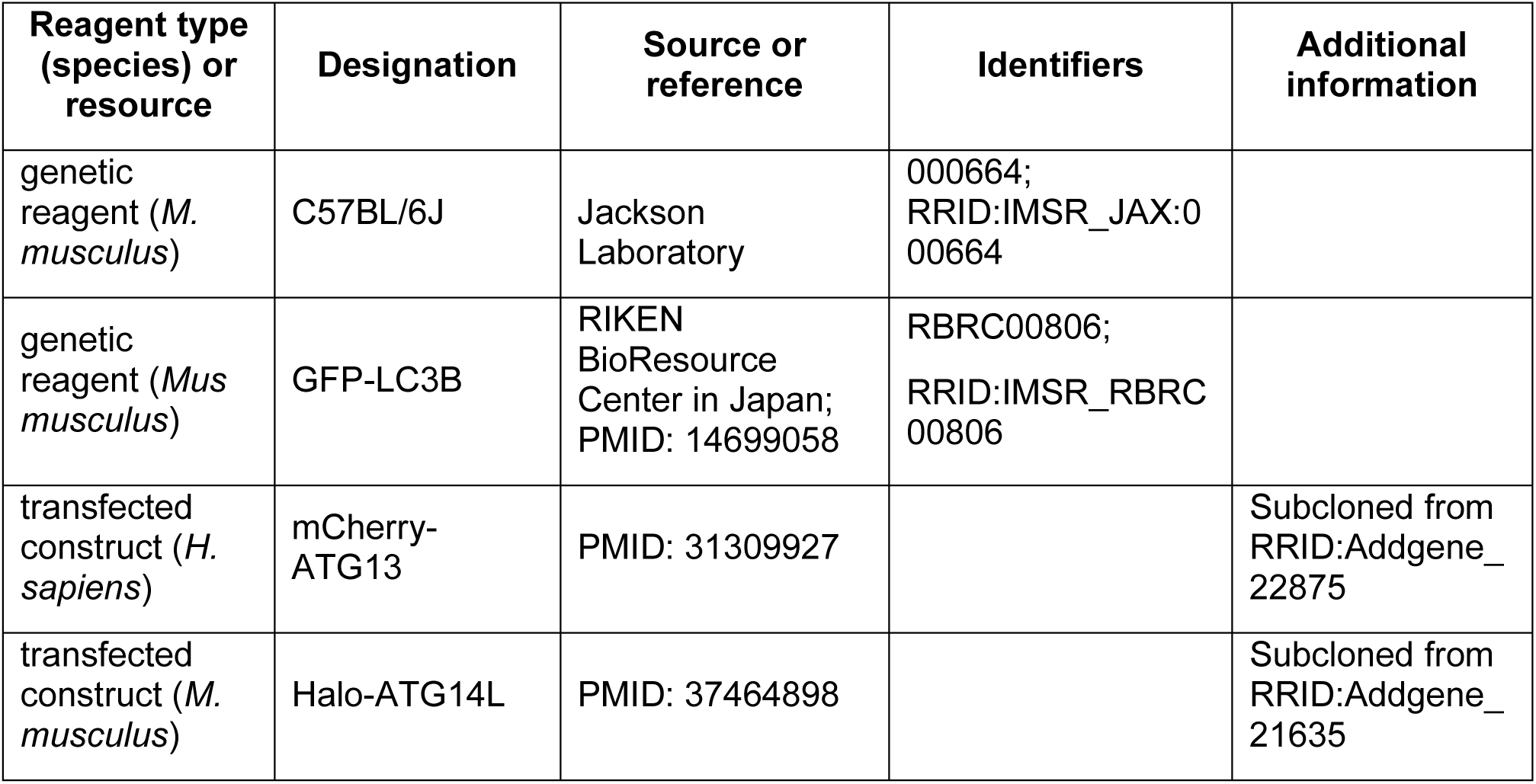
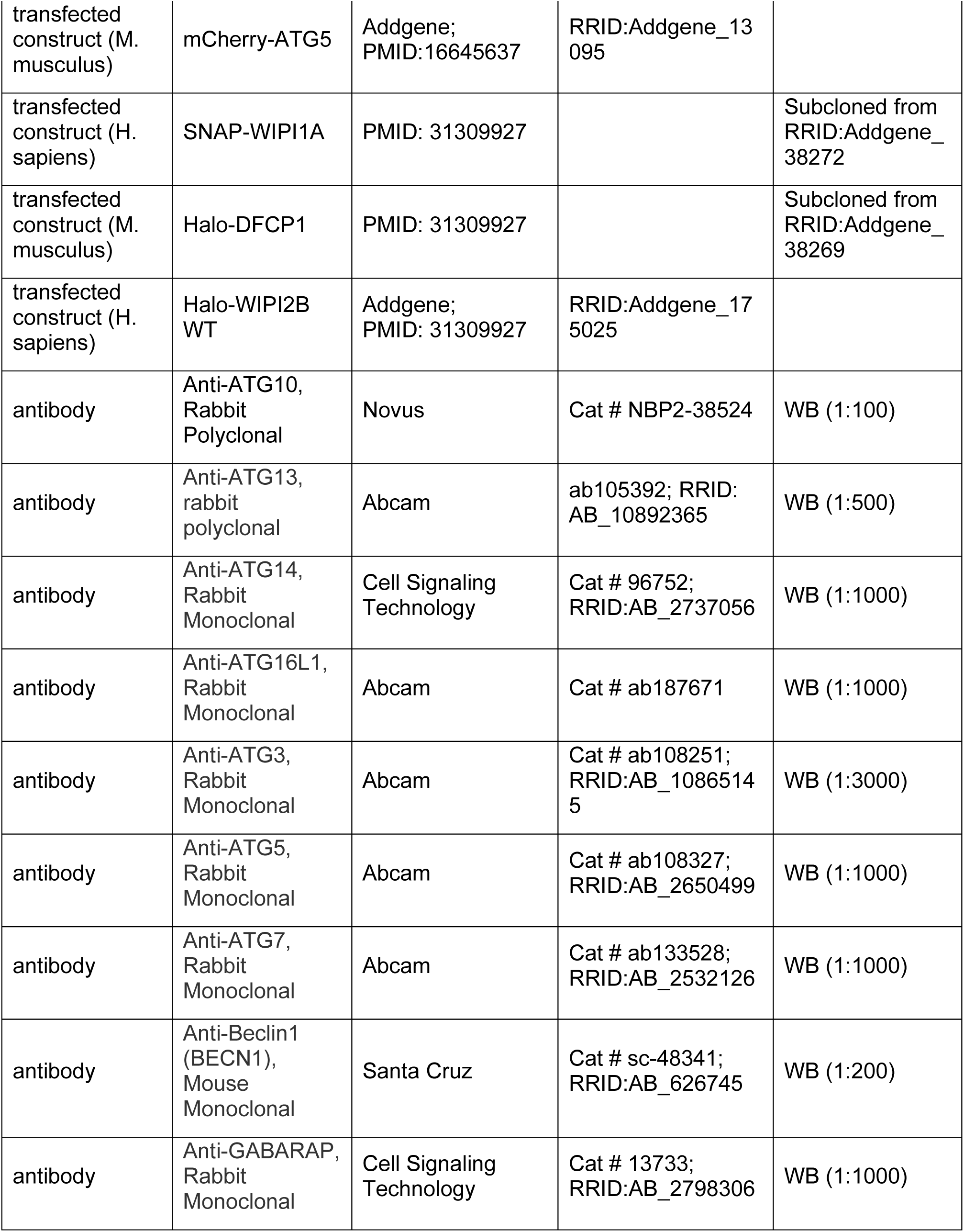
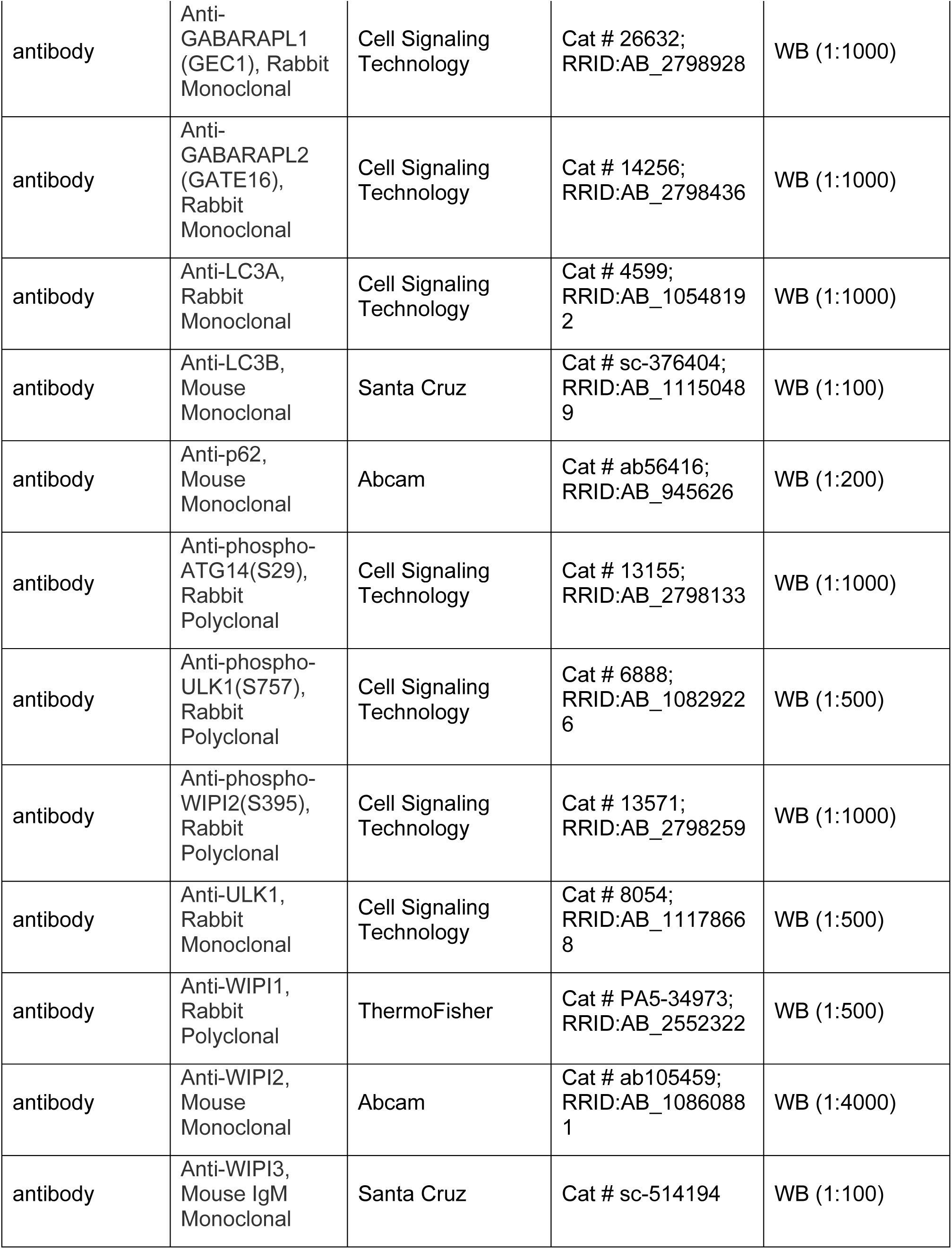
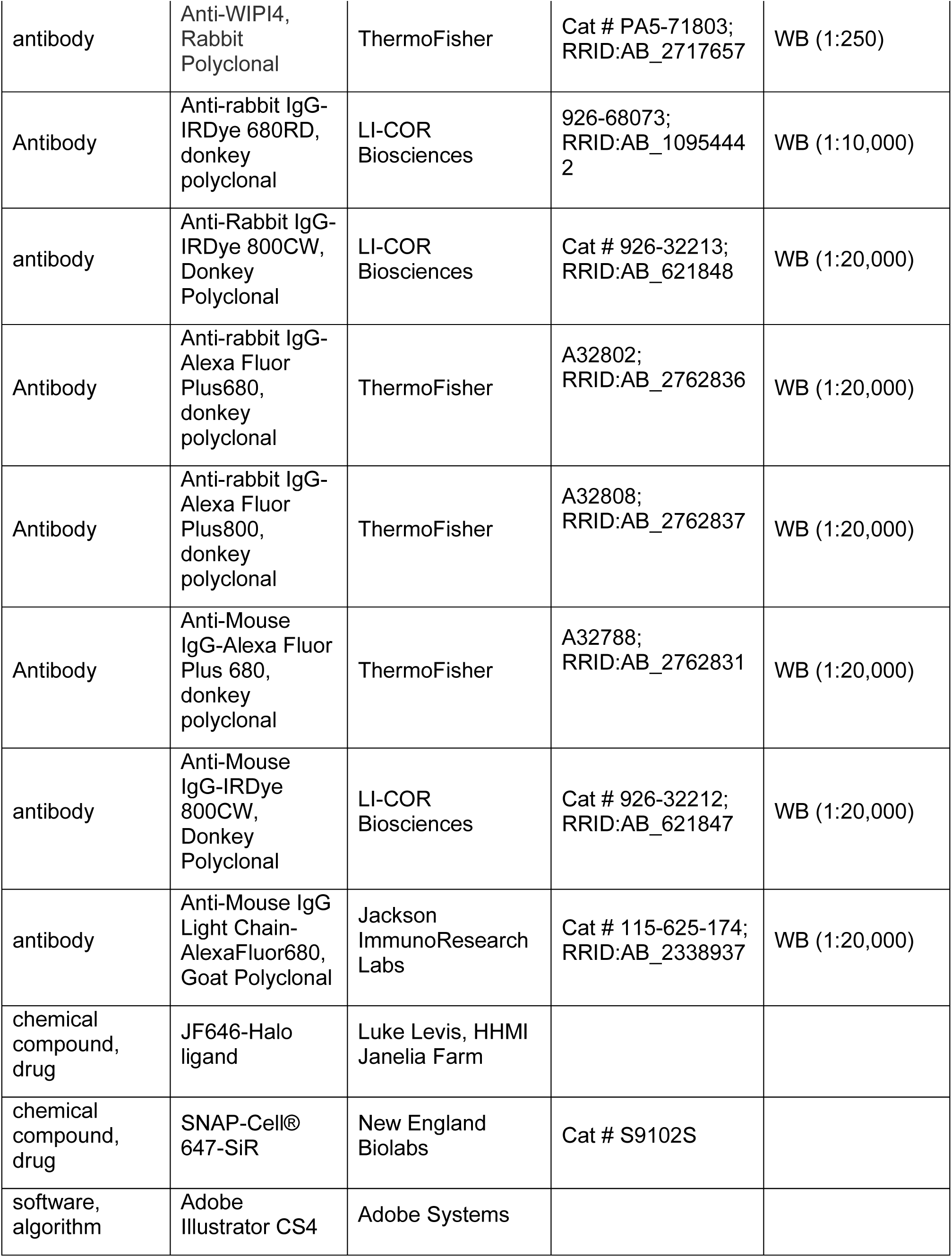
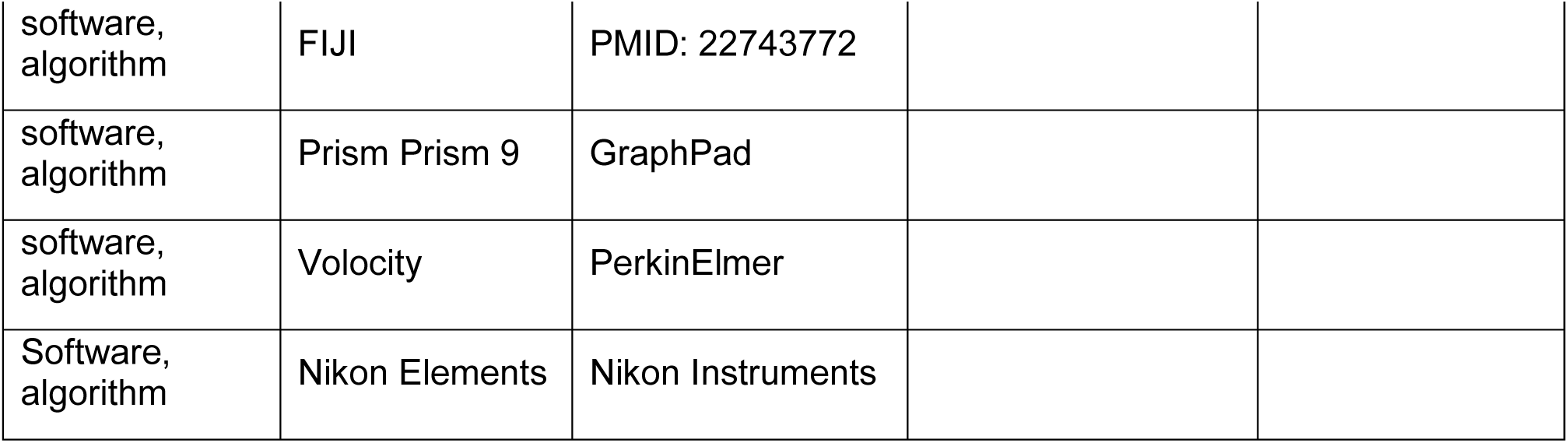
Key resources.

## ACKNOWLEDGEMENTS

The authors gratefully acknowledge members of the Stavoe and Waxham labs for comments on the manuscript.

