## Supplemental Files for "Sex does not Influence Neuronal Autophagy throughout Aging in Mice"

### SUPPLEMENTAL FIGURE LEGENDS

#### Figure S1. Induction complex protein levels.

**(A)** Immunoblots of DRG lysates from female and male young, young adult, aged, and advanced aged mice (n = 3 biological replicates shown for each sex). Total protein was used as a loading control (the normalization factor is indicated below each blot as a percentage). **(B)** Quantification of immunoblots in (A), normalized to total protein and represented as a fold change relative to the mean for lysates from young mice (mean ± sem). ns, not significant, by two-way ANOVA with Šídák's multiple comparisons test. **(C)** Immunoblots of brain lysates from female and male young, young adult, aged, and advanced aged mice (n = 3 biological replicates shown for each sex). Total protein was used as a loading control (the normalization factor is indicated below each blot as a percentage). **(D)** Quantification of immunoblots in (C), normalized to total protein and represented as a fold change relative to the mean for lysates from young mice (mean ± sem). ns, not significant; **p < 0.005 by two-way ANOVA with Šídák's multiple comparisons test.

#### Figure S2. Nucleation complex protein levels.

**(A)** Immunoblots of DRG lysates from female and male young, young adult, aged, and advanced aged mice (n = 3 biological replicates shown for each sex). Total protein was used as a loading control (the normalization factor is indicated below each blot as a percentage). **(B)** Quantification of immunoblots in (A), normalized to total protein and represented as a fold change relative to the mean for lysates from young mice (mean ± sem). ns, not significant by two-way ANOVA with Šídák's multiple comparisons test. **(C)** Immunoblots of brain lysates from female and male young, young adult, aged, and advanced aged mice (n = 3 biological replicates shown for each sex). Total protein was used as a loading control (the normalization factor is indicated below each blot as a percentage). **(D)** Quantification of immunoblots in (C), normalized to total protein and represented as a fold change relative to the mean for lysates from young mice (mean ± sem). ns, not significant; **p < 0.005 by two-way ANOVA with Šídák's multiple comparisons test. **(E)** Quantification of the rate of Halo-DFCP1 puncta formation in live-cell imaging of DRG neurons from female and male aged mice (median ± quartiles; n ≥ 29 neurons from three biological replicates of each sex). ns, not significant by unpaired t test (p = 0.89).

#### Figure S3. Elongation complex protein levels

**(A)** Immunoblots of DRG lysates from female and male young, young adult, aged, and advanced aged mice (n = 3 biological replicates shown for each sex). Total protein was used as a loading control (the normalization factor is indicated below each blot as a percentage). **(B-D)** Quantification of immunoblots in (A), normalized to total protein and represented as a fold change relative to the mean for lysates from young mice (mean ± sem). ns, not significant; *p < 0.05; **p < 0.005 by two-way ANOVA with Šídák's multiple comparisons test. **(E)** Immunoblots of brain lysates from female and male young, young adult, aged, and advanced aged mice (n = 3 biological replicates shown for each sex). Total protein was used as a loading control (the normalization factor is indicated below each blot as a percentage). **(F-H)** Quantification of immunoblots in (E), normalized to total protein and represented as a fold change relative to the mean for lysates from young mice (mean ± sem). ns, not significant; *p < 0.05 by two-way ANOVA with Šídák's multiple comparisons test.

**Figure S4. mATG8s and p62 protein levels (DRGs)**

**(A)** Immunoblots of DRG lysates from female and male young, young adult, aged, and advanced aged mice (n = 3 biological replicates shown for each sex). Total protein was used as a loading control (the normalization factor is indicated below each blot as a percentage). **(B-E)** Quantification of immunoblots in (A), normalized to total protein and represented as a fold change relative to the mean for lysates from young mice (mean ± sem). ns, not significant; *p < 0.05; **p < 0.005, ***p < 0.0005 by two-way ANOVA with Šídák's multiple comparisons test.

**Figure S5. mATG8s and p62 protein levels (brain)**

**(A)** Immunoblots of brain lysates from female and male young, young adult, aged, and advanced aged mice (n = 3 biological replicates shown for each sex). Total protein was used as a loading control (the normalization factor is indicated below each blot as a percentage). **(B-E)** Quantification of immunoblots in (A), normalized to total protein and represented as a fold change relative to the mean for lysates from young mice (mean ± sem). ns, not significant; *p < 0.05, by two-way ANOVA with Šídák's multiple comparisons test.

**Figure S6. WIPI3 protein levels**

**(A)** Immunoblots of DRG lysates from female and male young, young adult, aged, and advanced aged mice (n = 3 biological replicates shown for each sex). Total protein was used as a loading control (the normalization factor is indicated below each blot as a percentage). **(B)** Quantification of immunoblots in (A), normalized to total protein and represented as a fold change relative to the mean for lysates from young mice (mean ± sem). ns, not significant; *p < 0.05, by two-way ANOVA with Šídák's multiple comparisons test. **(C)** Immunoblots of brain lysates from female and male young, young adult, aged, and advanced aged mice (n = 3 biological replicates shown for each sex). Total protein was used as a loading control (the normalization factor is indicated below each blot as a percentage). **(D)** Quantification of immunoblots in (C), normalized to total protein and represented as a fold change relative to the mean for lysates from young mice (mean ± sem). ns, not significant by two-way ANOVA with Šídák's multiple comparisons test.

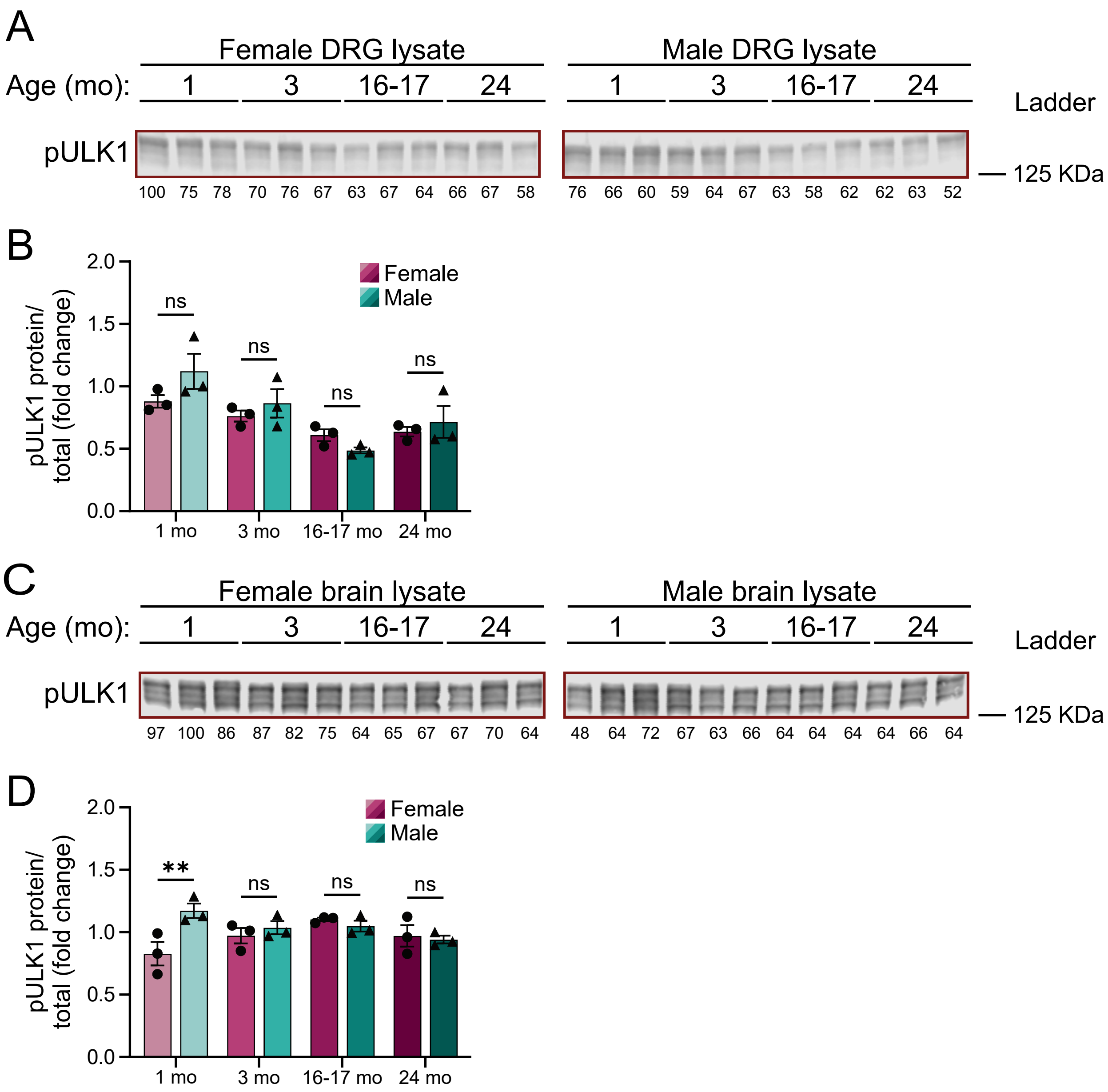
A
Female DRG lysate
Male DRG lysate
Age (mo): 1 3 16-17 24
1 3 16-17 24
Ladder
pULK1
125 KDa
100 75 78 70 76 67 63 67 64 66 67 58
76 66 60 59 64 67 63 58 62 62 63 52
B
pULK1 protein/ total (fold change)
0.0
0.5
1.0
1.5
2.0
ns
ns
ns
ns
Female
Male
1 mo
3 mo
16-17 mo
24 mo
C
Female brain lysate
Male brain lysate
Age (mo): 1 3 16-17 24
1 3 16-17 24
Ladder
pULK1
125 KDa
97 100 86 87 82 75 64 65 67 67 70 64
48 64 72 67 63 66 64 64 64 64 66 64
D
pULK1 protein/ total (fold change)
0.0
0.5
1.0
1.5
2.0
**
ns
ns
ns
Female
Male
1 mo
3 mo
16-17 mo
24 mo

FIGURE S1

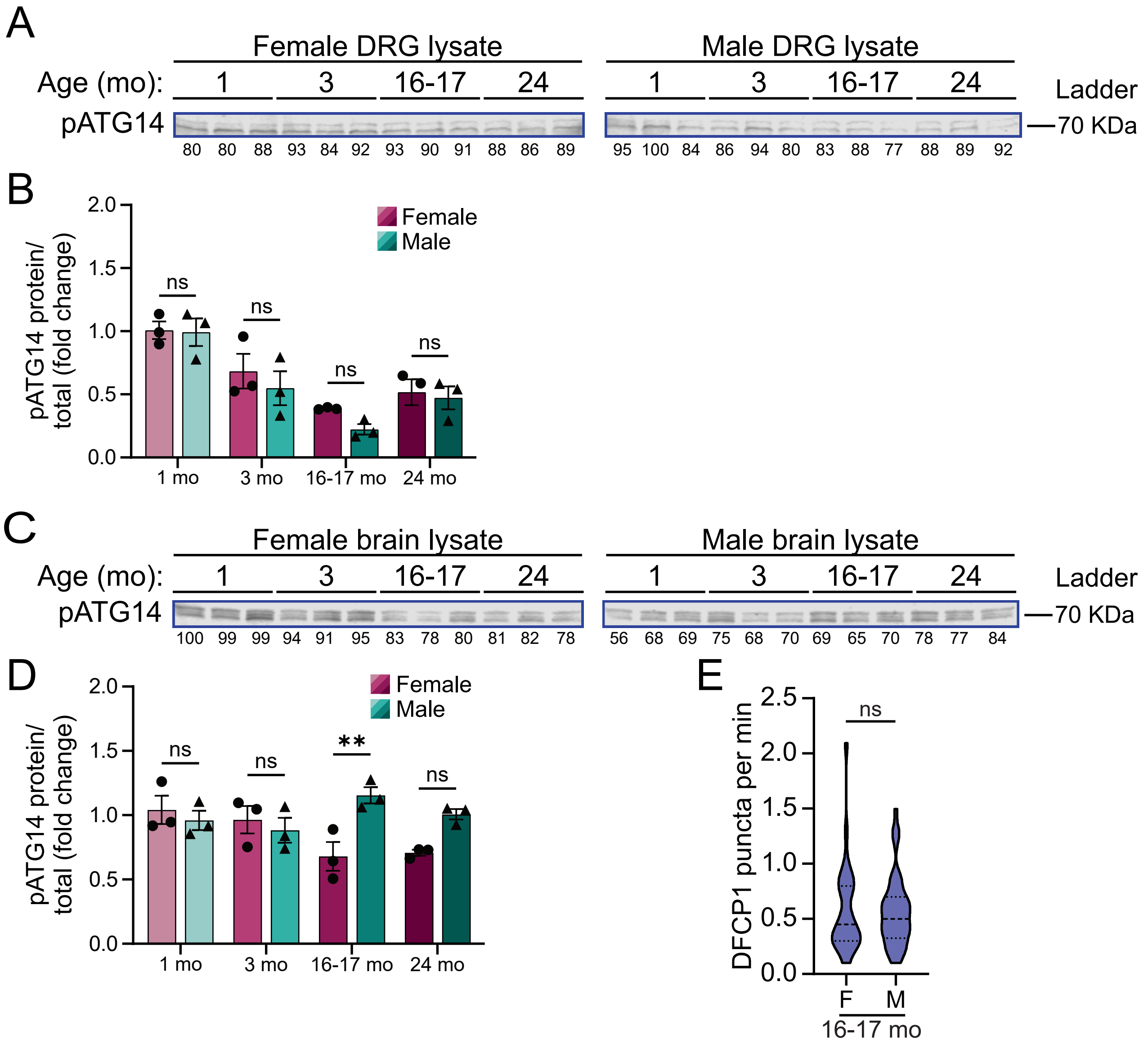
A
Female DRG lysate
Male DRG lysate
Age (mo): 1 3 16-17 24
Ladder
pATG14
—70 KDa
80 80 88 93 84 92 93 90 91 88 86 89
95 100 84 86 94 80 83 88 77 88 89 92
B
pATG14 protein/ total (fold change)
Female
Male
ns
1 mo 3 mo 16-17 mo 24 mo
C
Female brain lysate
Male brain lysate
Age (mo): 1 3 16-17 24
Ladder
pATG14
—70 KDa
100 99 99 94 91 95 83 78 80 81 82 78
56 68 69 75 68 70 69 65 70 78 77 84
D
pATG14 protein/ total (fold change)
Female
Male
ns
**
1 mo 3 mo 16-17 mo 24 mo
E
DFCP1 puncta per min
ns
F M
16-17 mo

FIGURE S2

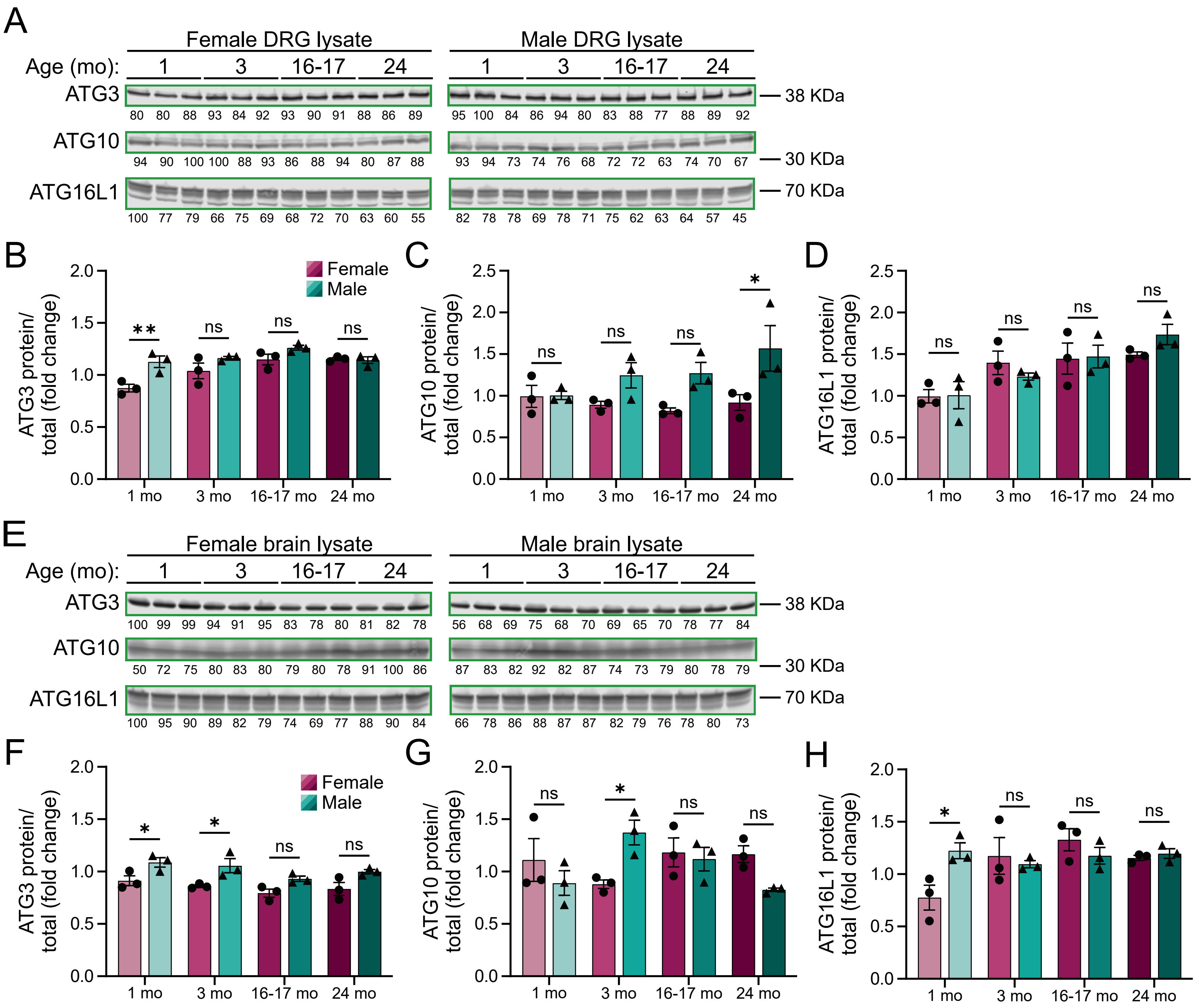

A
Female DRG lysate
Male DRG lysate
Age (mo): 1 3 16-17 24
ATG3
80 80 88 93 84 92 93 90 91 88 86 89
95 100 84 86 94 80 83 88 77 88 89 92
38 KDa
ATG10
94 90 100 100 88 93 86 88 94 80 87 88
93 94 73 74 76 68 72 72 63 74 70 67
30 KDa
ATG16L1
100 77 79 66 75 69 68 72 70 63 60 55
82 78 78 69 78 71 75 62 63 64 57 45
70 KDa
B
ATG3 protein/ total (fold change)
Female
Male
1 mo 3 mo 16-17 mo 24 mo
C
ATG10 protein/ total (fold change)
D
ATG16L1 protein/ total (fold change)
E
Female brain lysate
Male brain lysate
Age (mo): 1 3 16-17 24
ATG3
100 99 99 94 91 95 83 78 80 81 82 78
56 68 69 75 68 70 69 65 70 78 77 84
38 KDa
ATG10
50 72 75 80 83 80 79 80 78 91 100 86
87 83 82 92 82 87 74 73 79 80 78 79
30 KDa
ATG16L1
100 95 90 89 82 79 74 69 77 88 90 84
66 78 86 88 87 87 82 79 76 78 80 73
70 KDa
F
ATG3 protein/ total (fold change)
G
ATG10 protein/ total (fold change)
H
ATG16L1 protein/ total (fold change)


FIGURE S3

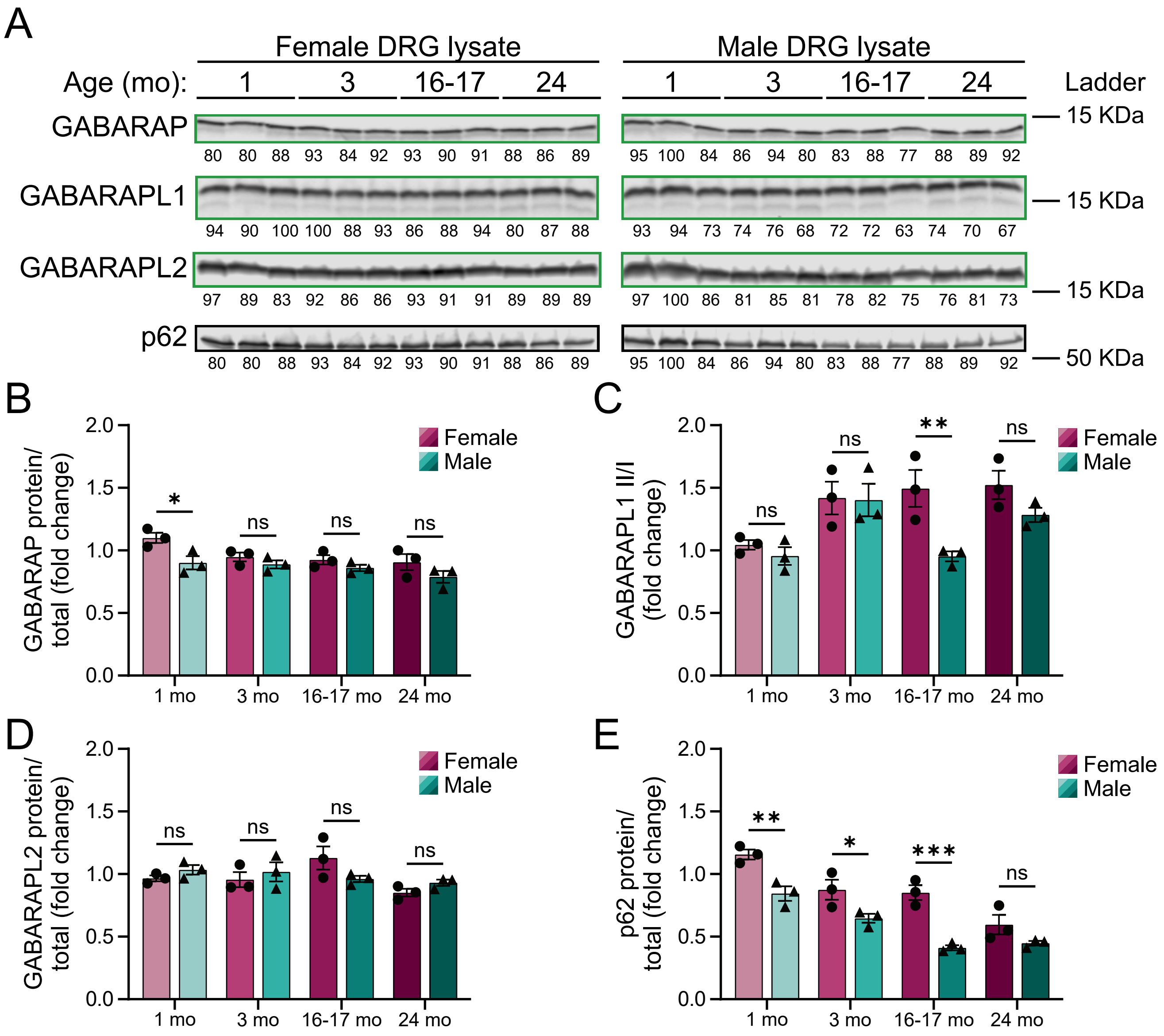
A
Female DRG lysate
Male DRG lysate
Age (mo):
1
3
16-17
24
Ladder
GABARAP
80 80 88 93 84 92 93 90 91 88 86 89
95 100 84 86 94 80 83 88 77 88 89 92
15 KDa
GABARAPL1
94 90 100 100 88 93 86 88 94 80 87 88
93 94 73 74 76 68 72 72 63 74 70 67
15 KDa
GABARAPL2
97 89 83 92 86 86 93 91 91 89 89 89
97 100 86 81 85 81 78 82 75 76 81 73
15 KDa
p62
80 80 88 93 84 92 93 90 91 88 86 89
95 100 84 86 94 80 83 88 77 88 89 92
50 KDa
B
GABARAP protein/ total (fold change)
Female
Male
1 mo
3 mo
16-17 mo
24 mo
C
GABARAPL1 II/I (fold change)
D
GABARAPL2 protein/ total (fold change)
E
p62 protein/ total (fold change)

FIGURE S4

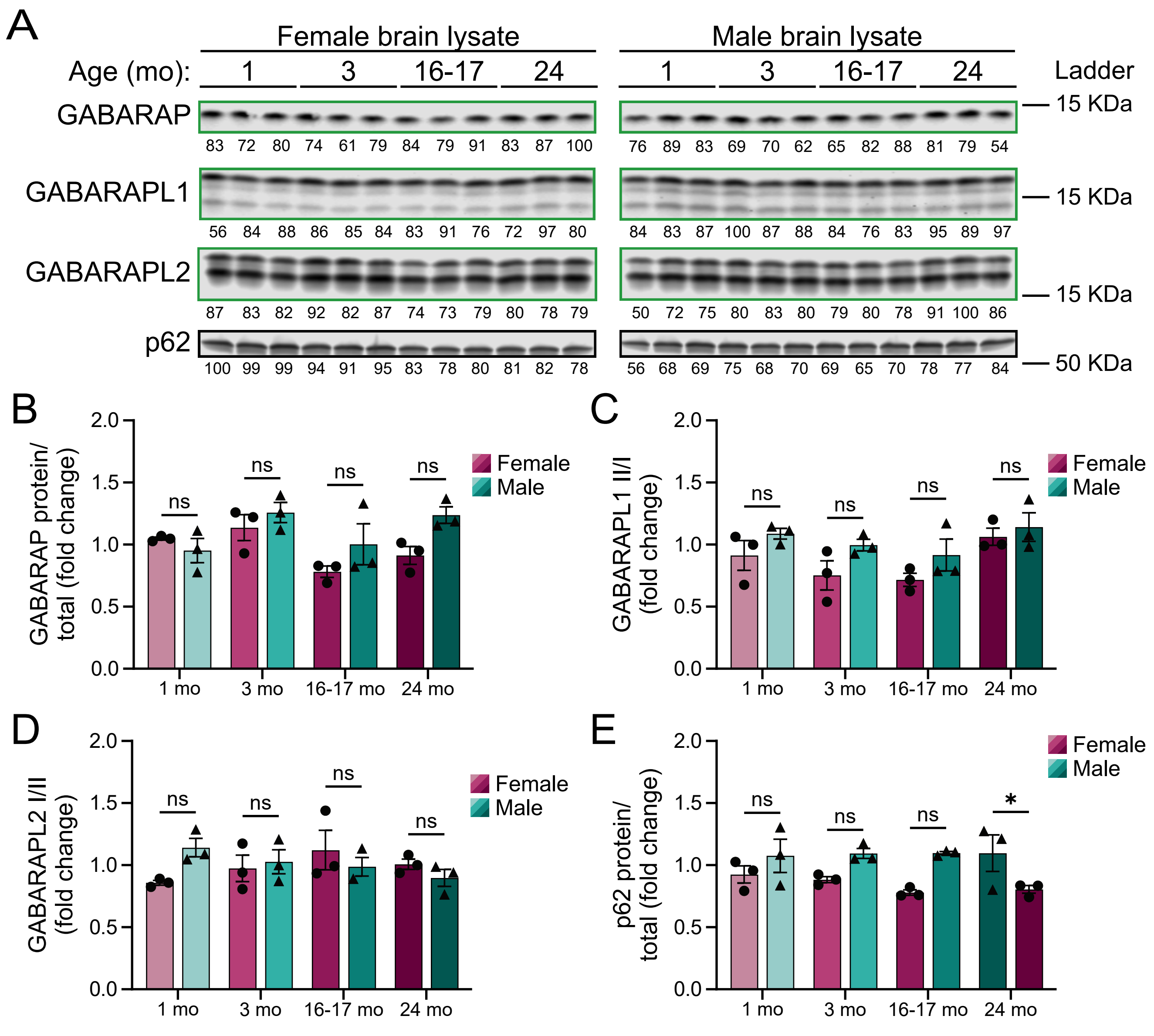

A
Female brain lysate
Male brain lysate
Age (mo): 1 3 16-17 24
Ladder
GABARAP
GABARAPL1
GABARAPL2
p62
15 KDa
50 KDa
83 72 80 74 61 79 84 79 91 83 87 100
76 89 83 69 70 62 65 82 88 81 79 54
56 84 88 86 85 84 83 91 76 72 97 80
84 83 87 100 87 88 84 76 83 95 89 97
87 83 82 92 82 87 74 73 79 80 78 79
50 72 75 80 83 80 79 80 78 91 100 86
100 99 99 94 91 95 83 78 80 81 82 78
56 68 69 75 68 70 69 65 70 78 77 84
B
GABARAP protein/ total (fold change)
C
GABARAPL1 II/I (fold change)
D
GABARAPL2 I/II (fold change)
E
p62 protein/ total (fold change)
ns
*
Female
Male
1 mo
3 mo
16-17 mo
24 mo


FIGURE S5

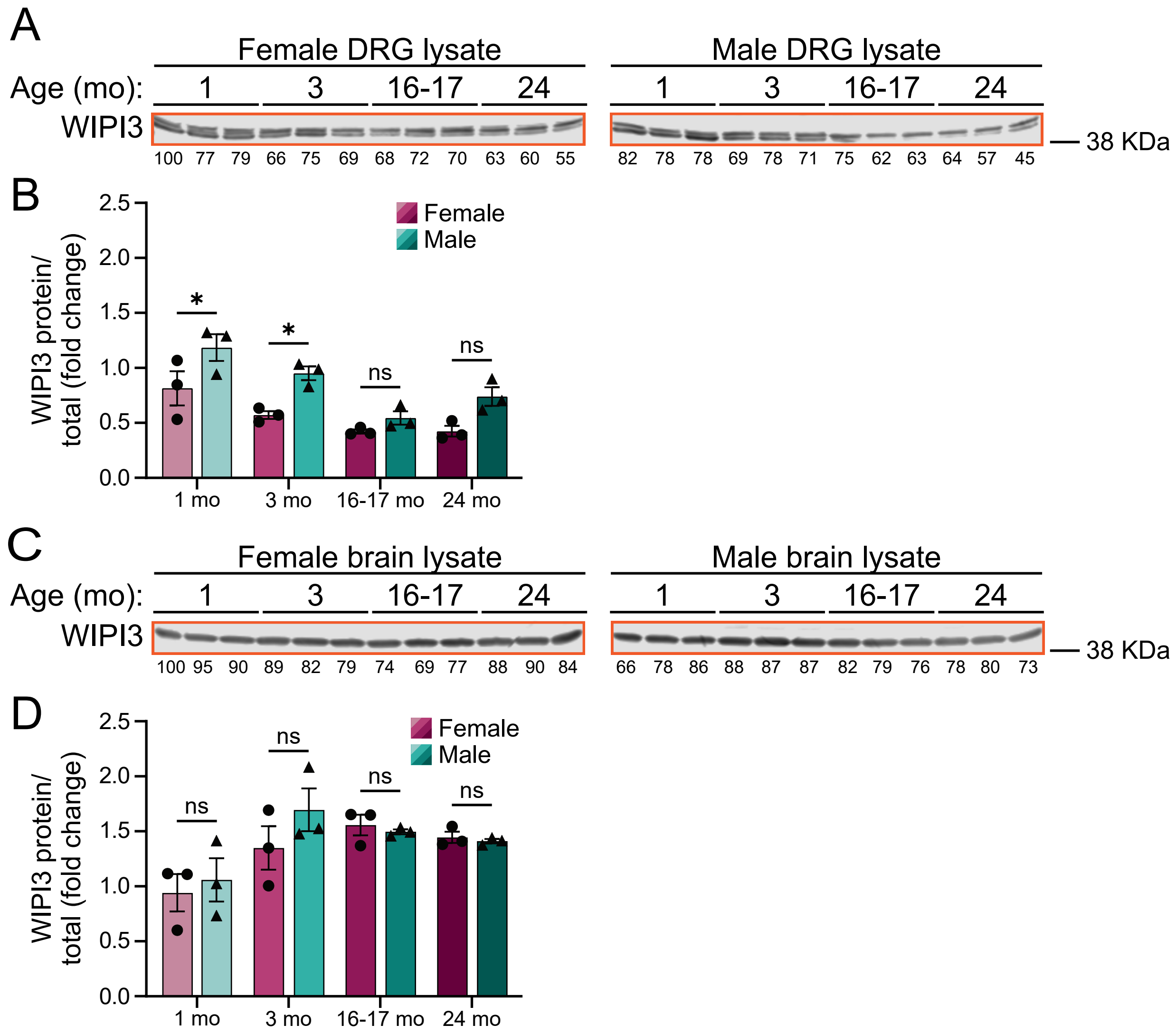
A
Female DRG lysate
Male DRG lysate
Age (mo): 1 3 16-17 24
WIPI3
38 KDa
100 77 79 66 75 69 68 72 70 63 60 55
82 78 78 69 78 71 75 62 63 64 57 45
B
WIPI3 protein/ total (fold change)
Female
Male
1 mo 3 mo 16-17 mo 24 mo
C
Female brain lysate
Male brain lysate
Age (mo): 1 3 16-17 24
WIPI3
38 KDa
100 95 90 89 82 79 74 69 77 88 90 84
66 78 86 88 87 87 82 79 76 78 80 73
D
WIPI3 protein/ total (fold change)
Female
Male
1 mo 3 mo 16-17 mo 24 mo

FIGURE S6
